# Mechanisms shaping the synergism of zeaxanthin and PsbS in photoprotective energy dissipation in the photosynthetic apparatus of plants

**DOI:** 10.1101/2021.01.10.426135

**Authors:** Renata Welc, Rafal Luchowski, Dariusz Kluczyk, Monika Zubik, Wojciech Grudzinski, Magdalena Maksim, Emilia Reszczynska, Karol Sowinski, Radosław Mazur, Artur Nosalewicz, Wieslaw I. Gruszecki

**Affiliations:** Department of Biophysics, Institute of Physics, Maria Curie-Sklodowska University, 20-031 Lublin, Poland; Institute of Agrophysics, Polish Academy of Sciences, 20-290 Lublin, Poland; Department of Plant Physiology and Biophysics, Institute of Biological Sciences, Maria Curie-Sklodowska University, 20-033 Lublin, Poland; Department of Metabolic Regulation, Institute of Biochemistry, Faculty of Biology, University of Warsaw, 02-096 Warsaw, Poland

**Author notes:** Corresponding Author: Wieslaw I. Gruszecki, Department of Biophysics, Institute of Physics, Maria Curie-Sklodowska University, 20-031 Lublin, Poland.

**Keywords:** photosynthesis, PsbS, zeaxanthin, xanthophyll cycle, photoprotection

## Abstract

Safe operation of photosynthesis is vital to plants and is ensured by the activity of numerous processes protecting chloroplasts against photo-damage. The harmless dissipation of excess excitation energy is believed to be the main photoprotective mechanism and is most effective with the simultaneous presence of PsbS protein and zeaxanthin, a xanthophyll accumulated in strong light as a result of the xanthophyll cycle activity. Here we address the problem of specific molecular mechanisms underlying the synergistic effect of zeaxanthin and PsbS. The experiments were conducted with *Arabidopsis thaliana*, the wild-type and the mutants lacking PsbS (*npq*4) and affected in the xanthophyll cycle (*npq*1), with the application of multiple molecular spectroscopy and imaging techniques. Research results lead to the conclusion that PsbS interferes with the formation of tightly packed aggregates of thylakoid membrane proteins, thus enabling the incorporation of xanthophyll cycle pigments into such structures. It was found that xanthophylls trapped within supramolecular structures, most likely in the interfacial protein region, determine their photophysical properties. The structures formed in the presence of violaxanthin are characterized by minimized dissipation of excitation energy. In contrast, the structures formed in the presence of zeaxanthin show enhanced excitation quenching, thus protecting the system against photo-damage.

## Introduction

Life on Earth is powered by the energy of sunlight, but utilization of this energy by living organisms is only possible thanks to the process of photosynthesis that converts the energy of electromagnetic radiation to the forms which can be directly used to drive biochemical reactions. Excess excitation is harmful to the photosynthetic apparatus as it can cause photo-oxidative damage (Ruban, 2016). During the biological evolution, photosynthetic organisms developed numerous photoprotective strategies that work at various organizational levels, including leaf movements, photo-translocation of chloroplasts in cells, or excessive excitation quenching in pigment-protein complexes, including the main plant antenna referred to as LHCII (Ruban et al., 2007; Croce and van Amerongen, 2014). The harmless dissipation of excessive excitation energy in the photosynthetic antenna systems is considered to be the primary photoprotective mechanism and can be monitored *in vivo* by the chlorophyll *a* (Chl *a*) fluorescence quenching (Ruban, 2016). Among the various phenomenological parameters describing Chl *a* fluorescence quenching, NPQ (non-photochemical quenching) turns out to be frequently used to monitor photoprotective energy dissipation in plant photosynthesis (Ruban and Wilson, 2020). It has been demonstrated that the most effective excitation quenching requires the simultaneous presence of both the PsbS protein (Li et al., 2000) and zeaxanthin (Zea) (Niyogi et al., 1998), a xanthophyll accumulated under high light conditions in consequence of de-epoxidation of violaxanthin (Vio) within the xanthophyll cycle (Jahns et al., 2009) (see Supplemental Figure 1). Moreover, overexpression of PsbS in plants results in enhanced light-induced excitation quenching (Steen et al., 2020). The significant quenching of Chl *a* excitations has been observed in reconstituted LHCII with the presence of Zea and PsbS, although neither PsbS nor Zea alone was sufficient to induce the same effect (Wilk et al., 2013). It has been demonstrated that the quenching of Chl *a* excitation associated with PsbS and Zea can occur independently (Holzwarth et al., 2009; Nilkens et al., 2010). In this respect, it is worth mentioning that PsbS-LHCII interaction in model membranes has been shown to induce highly effective chlorophyll excitation quenching in the pigment-protein complex (Pawlak et al., 2020). Regarding the role of Zea, there are two fundamentally different concepts proposed to explain photoprotective excitation quenching: one based on the direct contribution of this pigment to quenching of excessive excitations, and the other suggesting an indirect role of xanthophylls. It has been postulated that such quenching can be realized via the singlet-singlet excitation energy transfer between the Q_y_ energy level of Chl *a* and the S1 (^1^A_g_) level of Zea (Frank et al., 2000; Gruszecki et al., 2005). Alternatively, Chl *a* excitations can be quenched by the low-energy traps of the Zea cation radicals (Holt et al., 2005; Park et al., 2017). Recently, it has been demonstrated that both those mechanisms, the chlorophyll-carotenoid excitation energy transfer and the charge transfer can operate simultaneously as elements of regulatory activity in alga *Nannochloropsis oceanica* (Park et al., 2019). The singlet-singlet excitation energy transfer between the Q_y_ level of Chl *a* and the S1 level of carotenoids has been recently reported to operate in an intact trimeric structure of LHCII in the lipid environment (Son et al., 2020b). On the other hand, it has been shown that the replacement of Vio to Zea in the Vio-binding site of LHCII does not essentially influence the photophysical processes in the complex (Son et al., 2020a). As regarding an indirect role of the xanthophyll cycle in photoprotection, it was postulated that xanthophylls can be involved in the formation of supramolecular structures of the pigment-protein antenna complexes, characterized by quenching of potentially harmful, excessive excitation energy (Ruban et al., 1997; Gruszecki et al., 2006; Johnson et al., 2011; Xu et al., 2015; Welc et al., 2016; Zhou et al., 2020). Despite consensus on a photoprotective role of Zea synthesized within the xanthophyll cycle (Jahns and Holzwarth, 2012) detailed molecular mechanisms and photophysical processes responsible for excitation quenching involving xanthophylls are not fully understood. In particular, the synergism of Zea and PsbS in photoprotection seems to be one of the interesting and physiologically important problems awaiting clarification. In the present work, we address these problems in the study of intact leaves of *Arabidopsis thaliana* and isolated thylakoid membranes with the application of multiple molecular spectroscopy and imaging techniques specifically selected to address both the photophysical and structural issues.

## Results

### Accumulation of zeaxanthin in leaves

To obtain samples comprising different levels of Zea, leaves were illuminated for the same period at different light intensities. According to a detailed study on the kinetics of the xanthophyll cycle (Kress and Jahns, 2017), the time required for stabilization of Zea level in *A. thaliana* depends on an actual light intensity but also a genotype. For example, the equilibrium has not been observed even after 3 hours of illumination at 1800 µmol photons m^-2^s^-1^, in the case of *npq*4 plants (lacking PsbS). On the other hand, the same illumination for 2 hours of the WT leaves resulted in the stabilization of the Zea level at ca. 50 % of the xanthophyll cycle pool. In our work, we have arbitrarily selected a 30 min period for light exposure in experiments. Such a pre-illumination activated the de-epoxidation of Vio which resulted in the accumulation of Zea (see Figure 1A). As can be seen, the light intensity profile of Zea accumulation represents roughly a two-level dependency in the WT plants: Zea reaches a relatively low level (∼0.15) at light intensities below 600 µmol photons m^-2^s^-1^ and a level ∼0.3 at higher light intensities. Close Zea fractions for *A. thaliana* were observed for comparable light intensities and the same illumination period (30 min) (Kress and Jahns, 2017). Certain discrepancies can be attributed to the differences in plant growth conditions, in particular to photoperiod (long day versus short day). Importantly, a two-level light intensity profile of Zea accumulation, observed in the case of *A. thaliana* WT (Figure 1A) can also be observed in the case of other plant species (see Supplemental Figures 2 and 3). It is very likely that a two-level character of this dependency is an expression of light intensity-modulated availability of Vio to de-epoxidation (Jahns et al., 2009). In accordance with the original reports (Niyogi et al., 1998), light-induced accumulation of Zea has not been observed in the *npq*1 mutant lacking the de-epoxidase enzyme (Figure 1A). Interestingly, the Zea level was significantly higher in the case of the *npq*4 plants, particularly in the moderately high light intensity region (200-600 µmol photons m^-2^s^-1^). The direct explanation for this effect could be that the xanthophyll cycle is differently activated due to a different photosynthetic activity. The relatively low NPQ level in the *npq4* mutant (Figure 1B) implies a higher electron transfer rate through PSII and higher proton pumping resulting in a lower lumenal pH. On the other hand, no significant difference was observed in the PSII performance in the 200-600 µmol photons m^-2^ s^-1^ region (Figure 2A), where the *npq*4 shows a significant increase in Zea fraction (Figure 1A). This suggests that higher levels of Zea might be caused by a compensation type response. This could be realized at the level of gene expression of the xanthophyll cycle enzymes and/or via regulation of their activity. A similar compensation-type response consisting in accumulation of higher levels of Zea has been reported in the case of the *vte*1 genotype of *A. thaliana* lacking tocopherol (Havaux et al., 2005). Both tocopherol and Zea are efficient antioxidants with high potential in the protection of the lipid phase of the thylakoid membrane against oxidative damage. On the other hand, the Zea levels in both genotypes WT and *npq*4 are close at relatively high light intensities thus suggesting that other possible determinants also have to be considered. It may also not be excluded that the presence of PsbS decreases the availability of Vio for de-epoxidation via influencing the molecular organization of the antenna proteins. It has been proposed that PsbS may act as a “seeding center” for LHCII oligomerization in the thylakoid membranes (Sacharz et al., 2017). In this respect, it is worth mentioning that the enhanced availability of Vio for de-epoxidation has been observed in the case of the intermittent-light-grown plants lacking most of the antenna proteins that can potentially bind xanthophylls from the lipid phase of the thylakoid membrane (Jahns, 1995) (see also Supplemental Figure 3B). The results suggest that a plausible explanation of a two-level nature of the light intensity profile of Zea accumulation can be associated with the presence of two pools of Vio: one relatively freely available for de-epoxidation and another one, more tightly bound to the functional pigment-protein complexes, which becomes available for de-epoxidation under high light conditions (Jahns et al., 2009; Janik et al., 2016). The critical point of such a concept is a potential of PsbS to influence xanthophyll localization/organization in the thylakoid membrane. This problem is addressed below, in spectroscopic studies of intact leaves and isolated thylakoid membranes.

**Figure 1.**
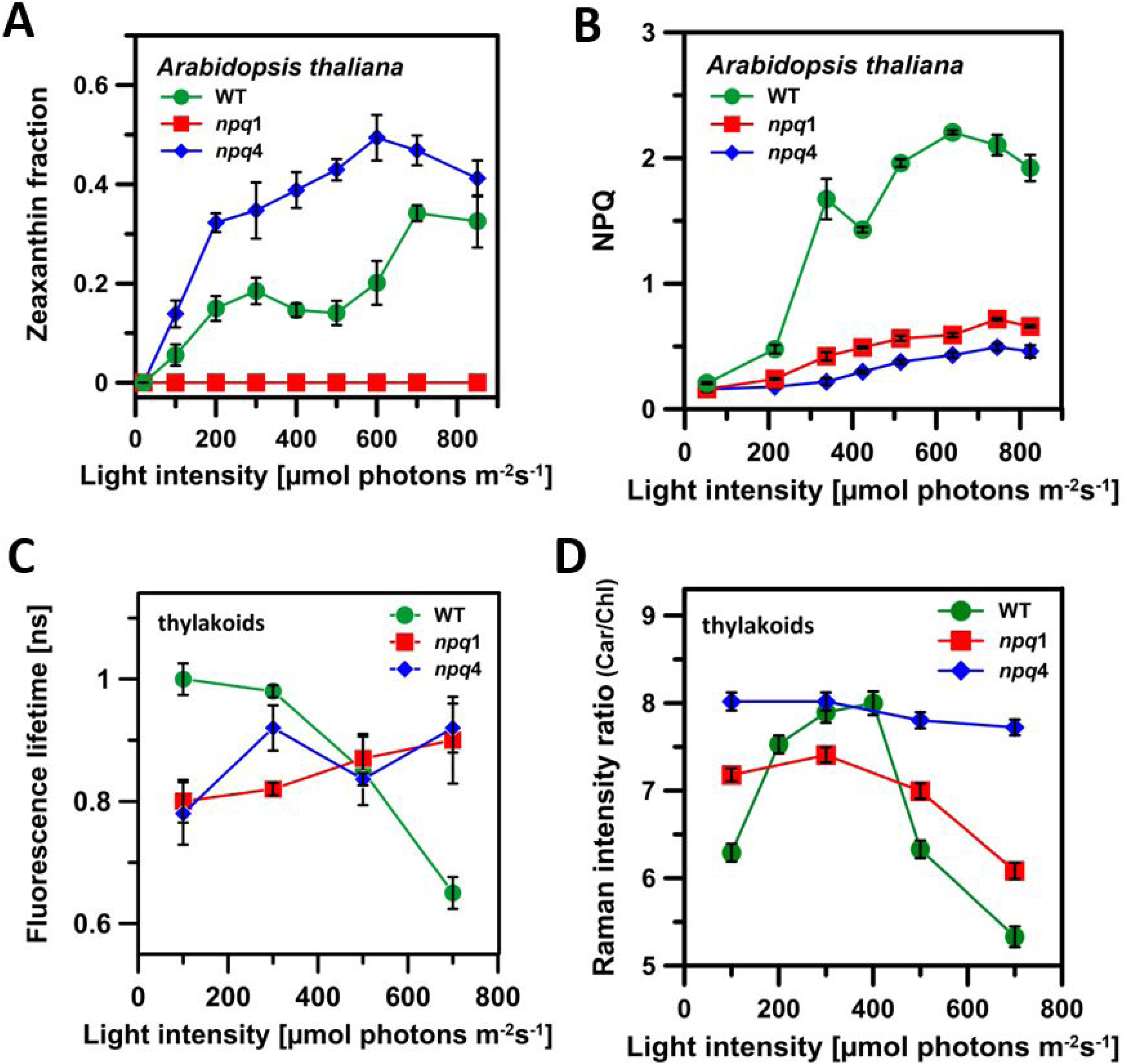
Light intensity dependencies of parameters determined in intact leaves and isolated thylakoid membranes. All the parameters were determined for the *A. thaliana* WT and the *npq*1 and *npq*4 genotypes, indicated. (A) Zeaxanthin level in leaves subjected to illumination for 30 min with white light at different intensities (represented in the abscissa axis). Zeaxanthin level was calculated as a fraction of the xanthophyll cycle pigments Zea/(Vio+Ant+Zea). The results represent mean from 6-12 experiments ± S.D. for the WT and *npq*4 samples and 3 experiments in the case of *npq*1. Each determination corresponding to one light intensity was made on a separate leaf from a different plant. (B) NPQ level determined in intact leaves. Experimental points represent NPQ values corresponding to 30 min of exposure to actinic light. The actinic light intensities are shown on the abscissa axis. The results represent mean from 4 experiments ± S.D. (C) Amplitude averaged fluorescence lifetime determined based on analyses of all photons collected in the course of imaging of multilayers formed with the thylakoid membranes. Prior to the thylakoid isolation, leaves were illuminated for 30 min with a white light of different intensity (represented in the abscissa axis). Exemplary images of multilayers are shown in Figure 2. Data represent the arithmetic mean from 4 experiments ± S.D. (D) Relative contribution of carotenoids to the Raman spectra of the thylakoid membranes isolated from leaves exposed to illumination for 30 min with a white light of different intensity (represented in the abscissa axis). The parameter (Car/Chl) was calculated as a ratio between the spectra integration results of the regions corresponding to carotenoids and chlorophylls (see Supplemental Figure 4). Experimental points represent the arithmetic mean from 120 spectra (the spectra from 30 pixels from 4 different samples) ± S.D. The data of the four panels were obtained from leaves and thylakoid preparations from the same batch. Light source and illumination conditions were identical for all the experiments except in the case of determination of NPQ (panel B). In such a case, actinic light was composed of a mixture of blue and red light of LED.

**Figure 2.**
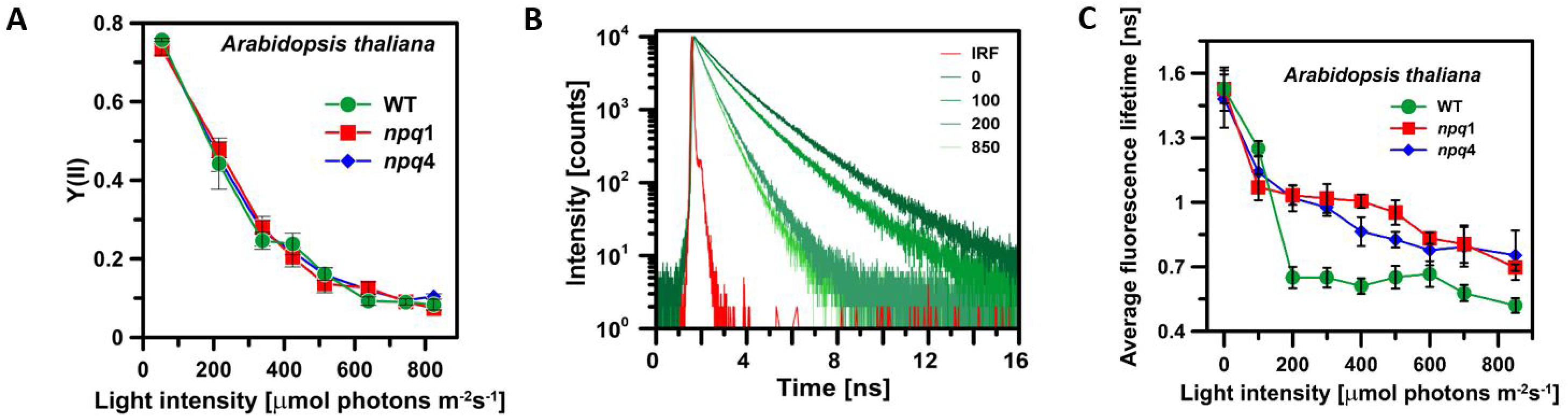
Chlorophyll fluorescence analysis in leaves exposed to different light intensities. (A) The dependency of the efficiency of photochemical reactions in Photosystem II (Y(II), based on modulated fluorescence analysis, in a function of intensity of white light used for preillumination (for 30 min) of *A. thaliana* leaves: WT, *npq*1 and *npq*4. The results represent mean from 4 experiments ± S.D. (B) Exemplary chlorophyll *a* fluorescence decay kinetics in intact *A. thaliana* WT leaves. Leaves were preilluminated for 30 min at different intensities of white light, indicated (in µmol photons m^-2^ s^-1^). The instrument response function (IRF) is also shown. (C) Intensity averaged fluorescence lifetime of chlorophyll *a* in intact leaves of *A. thaliana* determined after 30 min of illumination with white light at different intensity. Data represents the arithmetic mean from 6-9 experiments ± S.D.

**Figure 3.**
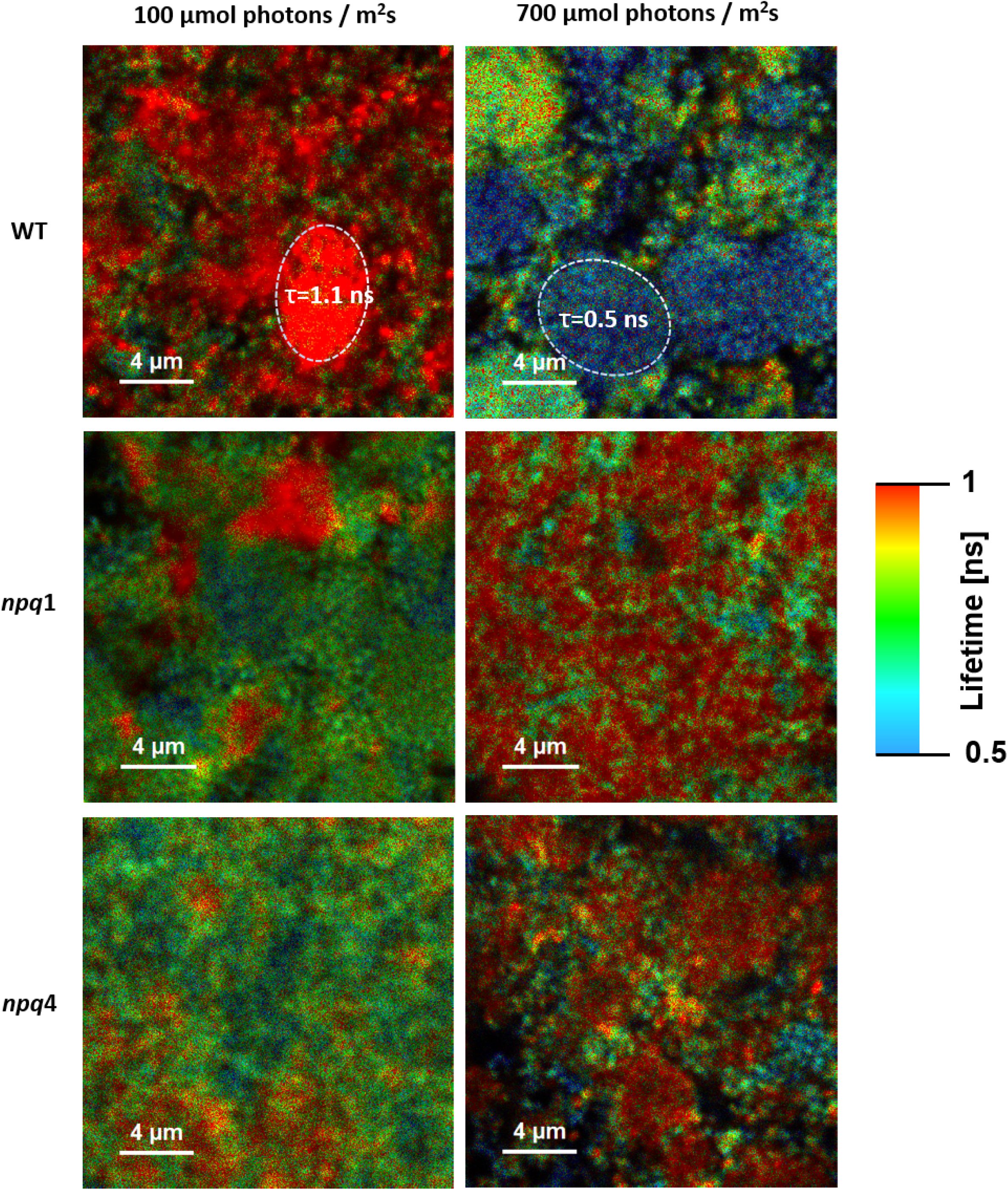
FLIM images of the multilayers formed of the thylakoid membranes. Thylakoid membranes were isolated from leaves of *A. thaliana*, three genotypes: WT, *npq*1 and *npq*4. Prior to isolation leaves were illuminated with white light at the intensity of 100 µmol photons m^-2^s^-1^ or 700 µmol photons m^-2^s^-1^ (indicated). Color codes represent Chl *a* fluorescence lifetimes (so-called fast FLIM). Amplitude averaged fluorescence lifetimes of two selected areas of the samples, representing the most “red” (long-lifetime) and the most “blue” (short-lifetime) regions, are presented in the top panels.

### Chlorophyll fluorescence quenching in leaves

As mentioned above, the photoprotective quenching of excessive excitation energy in the photosynthetic apparatus of plants can be analyzed within the framework of the non-photochemical quenching parameter (NPQ) of Chl *a* fluorescence in intact leaves (Krause and Weis, 1984; Ruban, 2016; Kress and Jahns, 2017). Figure 1B presents light intensity profiles of NPQ determined for *A. thaliana* WT as well as for the *npq*1 and *npq*4 mutants. As can be seen, the simultaneous presence of both Zea and PsbS protein is necessary for the photosynthetic apparatus to develop a significantly high photoprotective response, in accordance with the previous reports (Niyogi et al., 1998; Li et al., 2000; Steen et al., 2020). The levels of NPQ determined for different genotypes of *A. thaliana* in the present work are in agreement with the reported values corresponding to comparable light intensities (Kress and Jahns, 2017). The detailed analyzes of NPQ levels show that pronounced differences between WT plants and the *npq*1 and *npq*4 mutants are observed independently of actinic light intensity and a period of light exposure (Kress and Jahns, 2017). A measure of the synergistic effect of PsbS and Zea in photoprotective excitation quenching is the NPQ level higher by a factor of ∼ 2 in WT leaves compared to the sum of NPQ values determined for *npq*1 and *npq*4 plants, at high light intensities (above 500 µmol photons m^-2^ s^-1^, Figure 1B).

### Light-induced structural reorganization of the thylakoid membranes

Figure 3 presents fluorescence lifetime imaging microscopy (FLIM) images of ordered multilayers formed via deposition of thylakoid membranes at glass support. As can be seen, Chl *a* fluorescence lifetime, represented by color codes, depends on the pre-illumination history of plants from which the chloroplast membranes were isolated. FLIM analysis of the thylakoid membranes provides insight into molecular mechanisms associated with photoprotection due to the fact that Chl *a* fluorescence lifetime represents directly the regulatory quenching of singlet excitations (Sylak-Glassman et al., 2014; Chmeliov et al., 2016). Figure 1C presents average fluorescence lifetimes of Chl *a* in the thylakoid membranes isolated from plants exposed to different illumination. The average Chl *a* fluorescence lifetime values determined in the isolated thylakoid membranes (Figure 1C) correspond very well to the values reported for intact *A. thaliana* leaves, WT and both the *npq*1 and *npq*4 mutants (Sylak-Glassman et al., 2014). As might be expected, strong light resulted in the photoprotective excitation quenching, manifested by shortening of the fluorescence lifetimes (Figure 3, Figure 1C). Interestingly, the pronounced effect can be observed in the case of the thylakoid membranes isolated from *A. thaliana* WT plants but not in the case of the *npq*1 and *npq*4 genotypes. Such an observation directly corresponds to the results of the analysis of NPQ level in intact leaves (Figure 1B). Interestingly, a level of the average fluorescence lifetime determined for both the *npq*1 and *npq*4 mutants, following the 30 min of light exposure, is lower from the one determined for WT plants at low light intensities but the opposite trend is observed at strong light. This effect observed in the isolated thylakoid membranes suggests that simultaneous presence of both Zea and PsbS can act in the photosynthetic apparatus not exclusively to enhance quenching of excessive excitation energy under high light conditions but also protect against dissipation of excitation energy, highly undesirable under low light conditions. A similar tendency can be observed in the Chl *a* fluorescence lifetime analysis of intact *A. thaliana* leaves (Figure 2C), although the effect is not as pronounced as in the case of the isolated thylakoid membranes (Figure 1C). This can be explained by the activity and possible interplay of compensatory-type processes, which is also manifested by a similar PSII quantum yield in the leaves, despite the different level of excitation quenching (see Figures 2A and 1B).

The same type of samples of the oriented thylakoid multi-bilayers were analyzed with the application of the imaging technique based on resonance Raman spectroscopy. The results are presented in Figure 1D and Supplemental Figure 4. The thylakoid samples were scanned with the 457 nm laser line, being in resonance with both the carotenoid and chlorophyll pigments. This allowed analyzing the relative contribution of carotenoids to a Raman signal, with respect to chlorophylls, based on the spectral region representing the -C=C-stretching vibrations (Car/Chl, Supplemental Figure 4). Figure 1D presents the dependence of a relative contribution of carotenoids to Raman spectra on the intensity of light applied for the illumination of leaves before the thylakoid membrane isolation. Despite similar composition of carotenoids in the WT and *npq*4 leaves dark-adapted and exposed to low light (see Figure 1A and Supplemental Table 1) the Car/Chl ratio determined based on the resonance Raman study for the oriented thylakoid membranes differs considerably for the WT and *npq*4 samples (Figure 1D). On the other hand, very close Car/Chl values were determined in the case of the membranes isolated from WT and *npq*4 leaves preilluminated with the light intensity of 300 µmol photons m^-2^ s^-1^ (Figure 1D) despite the pronounced difference in the xanthophyll cycle pigment composition (Figure 1A). This implies that the Car/Chl parameter does not reflect directly the xanthophyll composition but rather can be dependent on protein-rearrangement-related changes, pigment molecular organization and in particular localization and orientation. In our opinion, it is highly probable that the differences in the Car/Chl ratio reflect the different orientation of xanthophylls with respect to the plane-oriented multilayer samples (Grudzinski et al., 2017). Raman imaging was carried out with the application of polarized laser light, which makes this technique particularly sensitive to the orientation of chromophores with respect to the surface of the sample owing to photo-selection (Grudzinski et al., 2017). Xanthophyll molecules oriented in the plane of imaging and parallel to the electric vector of laser light give rise to a strong, resonance-enhanced Raman signal, in contrast to the molecules oriented perpendicularly to the electric vector, which give rise only to a substantially less intensive non-resonant Raman scattering (a scheme explaining the idea of photo-selection by carotenoids embedded in lipid membranes is presented in Supplemental Figure 5). The relatively lower intensity of the resonance Raman signal of carotenoids in this particular model system is highly probable because most pigments are bound to the pigment-protein complexes in such a way that the long chromophore axes (dipole transitions) are oriented with acute angles to the axis normal to the plane of the multilayer thylakoid membranes (Grudzinski et al., 2017). Relatively higher resonance for carotenoids observed in the case of the *npq4* membranes (Figure 1D) strongly suggests that PsbS protein, absent in this particular mutant, accounts in the WT plants for the ordering of a certain fraction of carotenoid molecules within the thylakoid membranes in a fashion that their chromophores adopt a more vertical orientation with respect to the membrane plane: span the lipid bilayer (Grudzinski et al., 2017). Taking into consideration a relatively low number of PsbS molecules with respect to the other membrane proteins, one can expect its indirect role in the stabilization of carotenoid localization and orientation. It is very likely that this mechanism is associated with an effect of PsbS on the molecular organization of LHCII into aggregated structures (Sacharz et al., 2017). The increase in the Car/Chl ratio, accompanying the increase in light intensity (Figure 1D), can be understood in terms of a “liberation” of a certain pool of carotenoids from the ordering influence of PsbS. Interestingly, such an effect corresponds to the increase in the Zea level (Figure 1A) and therefore can be potentially linked with the process of making Vio available for de-epoxidation. The subsequent decrease in the Car/Chl ratio, observed in the light intensity range above 400 µmol photons m^-2^s^-1^, represents the higher level of Zea accumulation (Figure 1A, D). The fact that this process is associated with a decrease in the resonance Raman signal assigned to carotenoids suggests the activity of certain ordering mechanisms within the membrane, for example, associated with binding of newly synthesized Zea to the transmembrane proteins or trapping into protein supramolecular structures. Since such an effect is not observed in the case of the *npq*4 samples, it can be postulated that incorporation of xanthophylls into such structures is facilitated in the presence of PsbS. In this context, it is worth mentioning that the influence of PsbS on the organization of antenna proteins in the thylakoid membranes exposed to strong light has been reported (Sacharz et al., 2017). In the case of the *npq1* thylakoids, the changes in the Car/Chl ratio are similar to those observed in the case of the WT thylakoids (Figure 1D), although manifested to a much lesser extent. This suggests a lower susceptibility of epoxy-xanthophylls to a reorientation in the membrane, whether such stabilization is related to a pigment-protein interaction or the location and orientation of xanthophyll in the lipid phase of the membrane (see Supplemental Figure 6). It is also possible that the more planar orientation of a certain fraction of the membrane-bound carotenoids is associated with direct binding to the de-epoxidase enzyme (an increase in the Car/Chl ratio at low light intensities). The de-epoxidase belongs to a class of water-soluble lipocalin enzymes docking at the membrane surface and able to harbor the entire carotenoid molecule (Arnoux et al., 2009).

In order to analyze further possible relationship between PsbS and Zea we performed Raman spectroscopy analyses of intact leaves with the application of the 514 nm laser, being particularly in resonance with the xanthophylls with longer conjugated double bonds. Figure 4A presents the resonance Raman spectra recorded from intact *A. thaliana* leaves in the wavenumber range characteristic of carotenoids. The main band (referred to as ν_1_), centering in the 1526-1530 cm^-1^ spectral region, represents the -C=C-stretching vibrations in a polyene chain. The exact position of the band representing this vibration is sensitive to a length of the conjugated double bond system, molecular configuration and interaction to the environment (Mendes-Pinto et al., 2013; Sek et al., 2020). For this reason, the ν_1_ band in the Raman spectra recorded *in vivo* has a relatively large width, as it collectively represents carotenoids differing in localization and chemical structure, in particular in the number of conjugated double bonds in a polyene chain. Apart from Zea, the 514 nm laser is also in a strong resonance with Vio (Grudzinski et al., 2016b) and one of the molecules of lutein (Lut) (Ruban et al., 2001) in the protein environment of LHCII, due to the bathochromic shift of their absorption spectra. Figure 4B presents the spectral changes accompanying de-epoxidation of Vio to Zea resulting in the increase in the conjugation length from N=9 to N=11 (see Supplemental Figure 1) and is manifested by the spectral shift of the ν_1_ band towards lower wavenumbers (Ruban et al., 2001). Such a pronounced shift is visible both in the case of the WT and the *npq*4 plants but not in the case of the *npq*1 leaves. The higher amplitude of illumination-related spectral changes observed in the case of the *npq*4 plants as compared to the WT leaves corresponds to the higher level of Zea in the mutant. Figure 4C shows differences between the resonance Raman spectra of carotenoids in *A. thaliana* WT and *npq*1 and *npq4* mutants. The difference between WT and *npq*1 spectra might be associated to a certain degree with presence of a small fraction of antheraxanthin (Ant). On the other hand, a shift towards lower wavenumbers of the entire carotenoid spectrum recorded in the WT with respect to that one corresponding to the *npq*1 thylakoids suggests rather different molecular organization/binding properties of pigments in the mutant. Interestingly, very small differences can be observed between the WT and *npq*4 plants (Figure 4C). PsbS has been shown not to be a canonical pigment-binding protein (Fan et al., 2015). This suggests that if xanthophylls interact with the PsbS protein then with its surface, at the border with the lipid phase of the membrane, and in a nonspecific manner (Bonente et al., 2008). The relatively small magnitude of the spectral changes observed in Figure 4C leads to the conclusion that putative interaction of PsbS with xanthophylls is not responsible for the pronounced effects shown in Figure 1D. This implies that the effect of PsbS on xanthophyll orientation in the thylakoid membranes is rather realized indirectly, by influencing the molecular organization of pigment-protein complexes.

**Figure 4.**
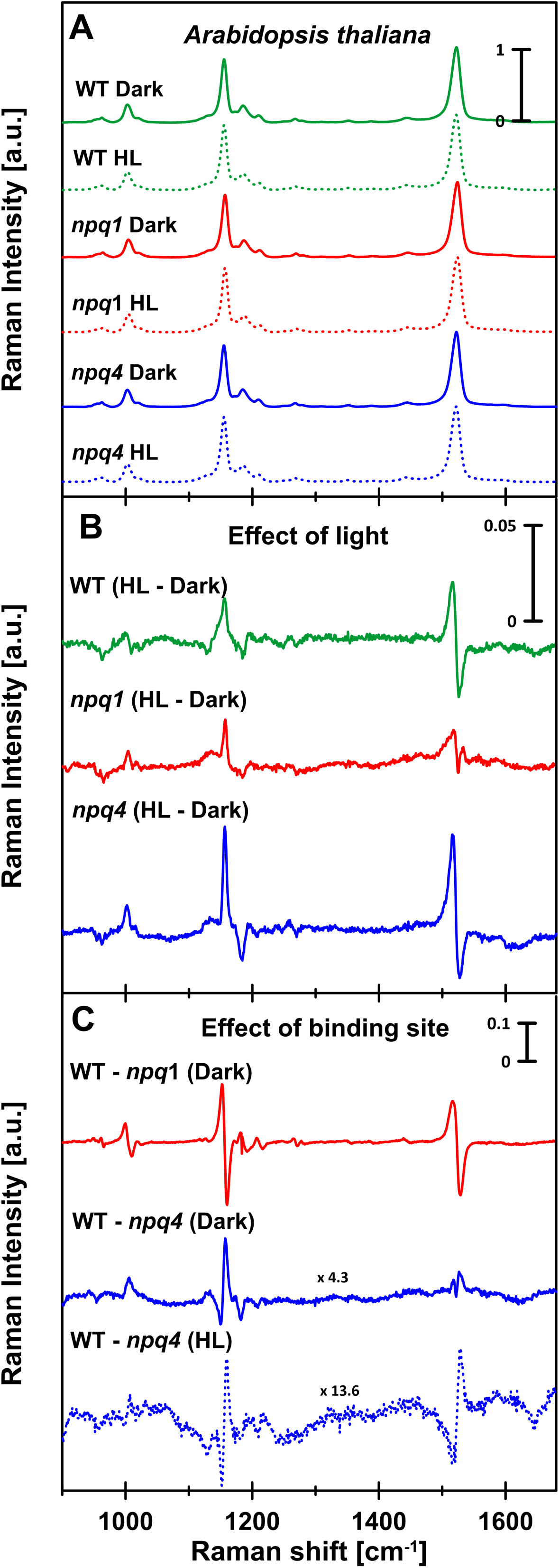
Resonance Raman spectra of leaves. Spectra recorded with 514 nm laser. (A) The spectra were recorded from leaves of three genotypes of *A. thaliana*: WT, *npq*1 and *npq*4, dark-adapted or pre-illuminated for 180 min with white light, the intensity of 1800 µmol photons m^-2^s^-1^ (marked either Dark or HL). Panels B and C present the difference spectra calculated based on the spectra presented in panel A. Panel B presents the effect of illumination and panel C presents the spectral effects associated with the absence of violaxanthin de-epoxidase (*npq*1) or PsbS protein (*npq*4). The spectra presented in panel A were normalized at the maximum. Each spectrum represents an average calculated from 1300 individual spectra recorded in a leaf area of 130 x 180 µm. The relative xanthophyll cycle pool composition was: WT Dark Vio 0.92, Ant 0.08, Zea 0.0; WT HL Vio 0.31, Ant 0.12, Zea 0.57; *npq*1 Dark Vio 1.0, Ant 0.0, Zea 0.0, *npq*1 HL Vio 1.0, Ant 0.0, Zea 0.0, *npq*4 Dark Vio 0.93, Ant 0.07, Zea 0.0, *npq*4 HL Vio 0.27, Ant 0.20, Zea 0.53.

Possible effect of PsbS and Zea on molecular organization of the thylakoid membrane proteins was analyzed in the present study with the application of FTIR spectroscopy. Figure 5 and Supplemental Figure 7 present the IR absorption spectra of the oriented multibilayers formed of the thylakoid membranes (such as those imaged with the application of the FLIM technique, Figure 3). Analysis of the amide I band (1600-1700 cm^-1^) gives an insight into the secondary structure of a protein, but also its molecular organization (Tamm and Tatulian, 1997; Zhou et al., 2020). The spectra recorded show a contribution from the components in the region of 1650 cm^-1^, which can be assigned to α-helical segments of the transmembrane functional proteins embedded in the thylakoid membranes and the region of 1630 cm^-1^, which can be assigned to pseudo-β-structures formed due to the formation of protein molecular assemblies, as documented in the case of LHCII in different model systems (Gruszecki et al., 2006; Janik et al., 2013; Zhou et al., 2020). Interestingly, the intensity of the spectral bands representing α-helical structures (1648-1660 cm^-1^) and turns and loops (1660-1685 cm^-1^), relative to the band representing aggregated forms of the membrane proteins (1620-1630 cm^-1^) (Tamm and Tatulian, 1997) is higher at low light in the case of WT plants than in the case of *npq*1 and *npq*4. As can be seen, illumination shifts the equilibrium towards protein aggregated structures in the case of the WT plants but has a little effect in the case of both *npq*1 and *npq*4 mutants (Figure 5, Supplemental Figure 7). Such an observation implies that illumination results in the formation of molecular assemblies of certain thylakoid membrane proteins, formed with the involvement of both Zea and PsbS. Importantly, the formation of the supramolecular domains by the LHCII proteins in the thylakoid membranes and their light-dependent reorganization was documented with the application of the freeze-fracture electron microscopy (Johnson et al., 2011).

**Figure 5.**
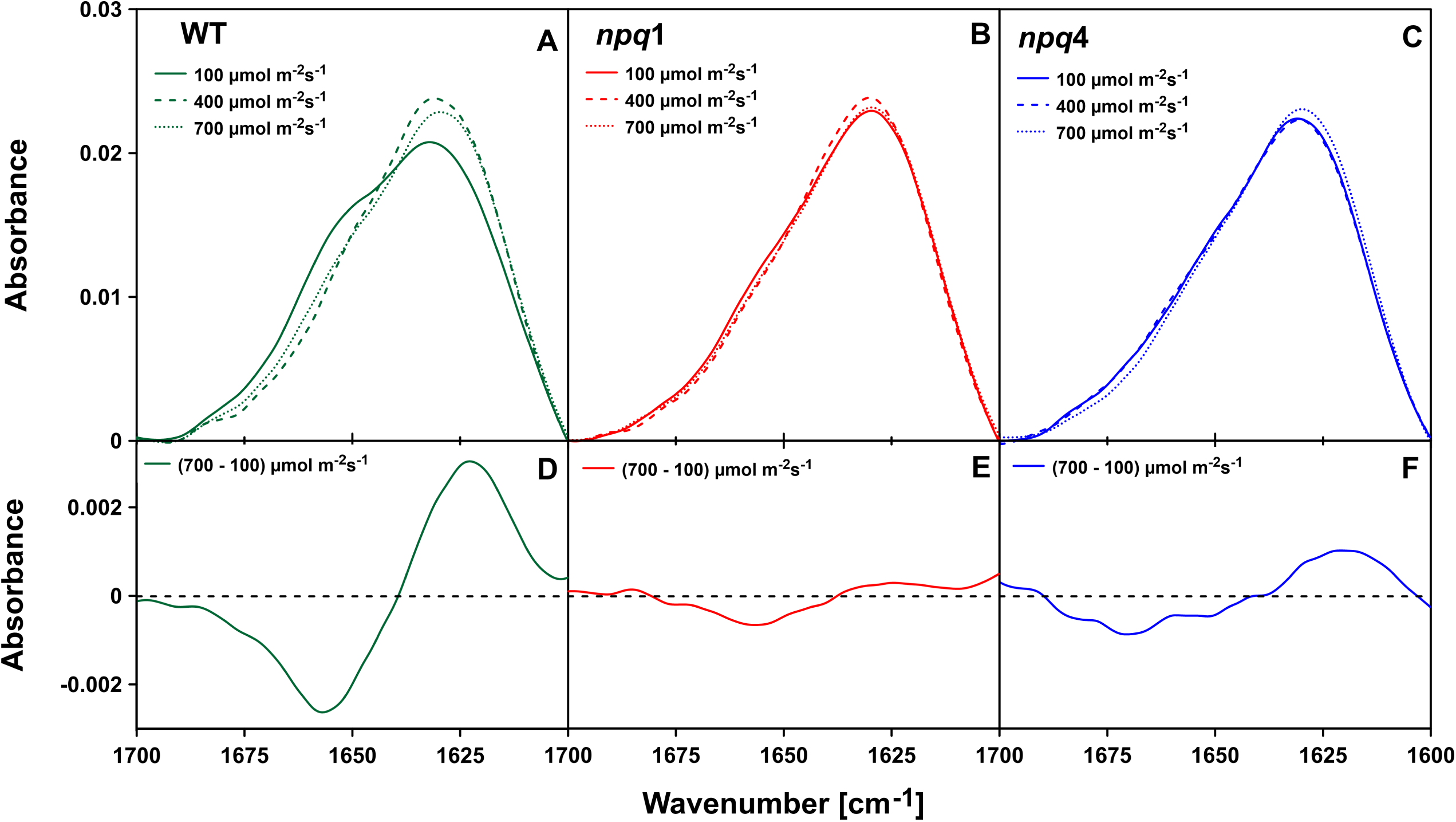
FTIR spectra of the thylakoid membranes. The infrared absorption spectra were recorded from the thylakoid membranes of three genotypes of *A. thaliana*: WT, *npq*1 and *npq*4, isolated from leaves pre-illuminated for 30 min with a white light of different intensities (indicated). Panels A, B and C present the original spectra and the panels D, E and F present the difference spectra. The original, surface-normalized spectra are presented in the amide I region (1600-1700 cm^-1^). The samples were dark-adapted under argon for 15 min before and during the measurements. Each spectrum in panels A, B and C represents a different experiment. An additional set of results is presented in Supplemental Figure 7.

Molecular organization of the photosynthetic pigment-protein complexes in the thylakoid membranes can be also analyzed based on the comparison of the circular dichroism (CD) spectra recorded from the thylakoid samples isolated from different genotypes of *A. thaliana* (Figure 6, Supplemental Figure 8). In general, the spectra recorded are characterized by the dominant bands at 439 nm (-), 462 nm (-), 507 nm (+), 653 nm (-) and 690 nm (+), in accordance with the previous reports (Toth et al., 2016). The CD bands in the spectra of the thylakoid membranes originate primarily from the excitonic and psi-type interactions between pigments bound to the functional pigment-protein complexes which are involved in the formation of supramolecular structures (Toth et al., 2016). Importantly, the CD spectra recorded from the *npq*4 samples differ significantly from the spectra recorded for the WT and *npq*1 thylakoids. At the absence of PsbS, the CD signal is stronger in the spectral regions representing both Chl *b* (653 nm, 462 nm) and carotenoids (507 nm), despite normalization at the maximum of the band representing Chl *a* (690 nm). This implies that PsbS interferes with the formation of ordered molecular assemblies of the antenna proteins accounting for the psi-type and/or excitonic interactions between the photosynthetic pigments.

**Figure 6.**
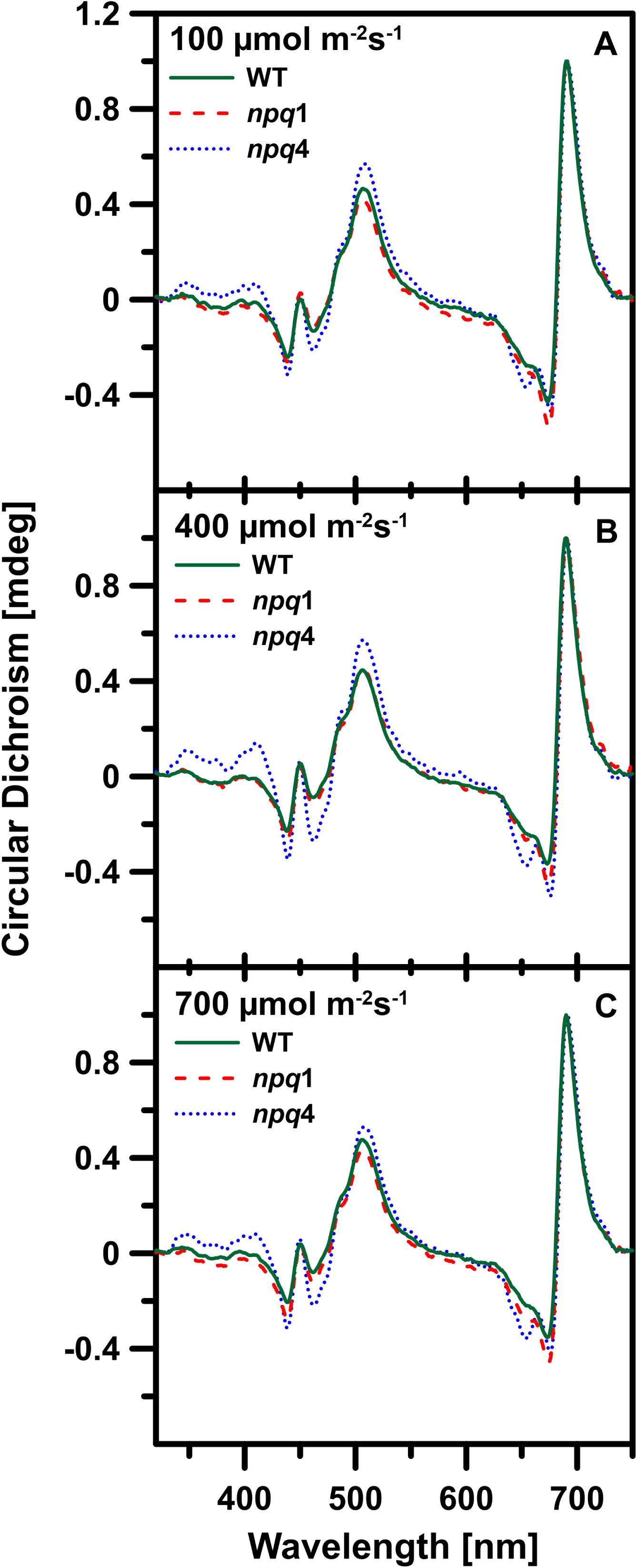
CD spectra of thylakoid membranes. The circular dichroism spectra were recorded from the thylakoid membranes of three genotypes of *A. thaliana*: WT, *npq*1 and *npq*4, isolated from leaves pre-illuminated for 30 min with a white light of different intensities (indicated). The samples were dark-adapted for 15 min before the measurements. The spectra were normalized at the maximum. Each spectrum represents a different experiment. An additional set of results is presented in Supplemental Figure 8.

## DISCUSSION

Analysis of light intensity profiles of Zea synthesis within the xanthophyll cycle shows a two-level character of this dependency (Figure 1A, Supplemental Figures 2, 3). Factors that are directly involved in governing the levels of Vio and Zea are activities of the key enzymes of the xanthophyll cycle: Vio de-epoxidase (VDE) and Zea epoxidase (ZEP). In principle, the activity of VDE is controlled by light intensity, indirectly, via the dependence on pH of the thylakoid lumen (Jahns et al., 2009). On the other hand, it has been demonstrated that also ZEP is light-controlled, indirectly, via the redox system associated with the NADPH thioredoxin reductase C (Naranjo et al., 2016). It is very likely that differences in the activity of VDE and ZEP at different light intensities can account for the variations in the level of the xanthophyll cycle pigments observed. Another factor that can be involved in a control of xanthophyll interconversion is the mechanism referred to as availability of Vio to de-epoxidation (Siefermann and Yamamoto, 1974; Siefermann-Harms, 1984; Wehner et al., 2006; Jahns et al., 2009). It is possible that the lower level of Zea (at light intensities below 600 µmol photons m^-2^s^-1^) is associated with de-epoxidation of the Vio pool less tightly bound to the pigment-protein complexes and therefore available more freely for the enzymatic de-epoxidation in the thylakoid membranes (Latowski et al., 2004; Jahns et al., 2009). The appearance of Vio pools characterized by different levels of availability for de-epoxidation, one relatively easily accessible, in the lipid phase of the thylakoid membrane and another one comprising xanthophylls which are originally protein-bound, has been postulated and observed in several previous studies (Hartel et al., 1996; Wehner et al., 2006; Janik et al., 2016). It has been even proposed, based on the results of the research and the literature analysis, that a certain fraction of Vio, not bound to the pigment-protein complexes, can be directly present in the lipid phase of the membranes (Janik et al., 2016; Welc et al., 2016). In our opinion, the higher level of Zea, observed at elevated light intensities (above ∼500 µmol photons m^-2^s^-1^) can be associated with the “liberation” of a new Vio pool, very likely trapped within the trimeric LHCII. According to the results of the FTIR analyses presented here (Figure 5, Supplemental Figure 7), a significant reorganization of the membrane proteins takes place at such light intensities, which are associated with the xanthophyll reorientation within the membrane, manifested by Raman spectroscopy (Figure 1D). In our opinion, the xanthophyll reorientation postulated can be associated with the process of making Vio available for enzymatic de-epoxidation. Photo-isomerization of Vio from the all-*trans* to 9-*cis* or 13-*cis* molecular configuration (Grudzinski et al., 2001) may be considered as a mechanism potentially involved in this process. On the other hand, a highly efficient light-driven back reaction to the all-*trans* molecular configuration is observed, even at very low light intensities (Sek et al., 2020). This mechanism can be considered a critical one for an effective de-epoxidation owing to the fact that Vio in the molecular configuration all-*trans* is the specific substrate for the VDE enzyme (Yamamoto and Higashi, 1978). The higher level of Zea accumulation, observed at light intensities above ∼500 µmol photons m^-2^ s^-1^ (Figure 1A, Supplemental Figure 2A), can be associated with the process of “liberation” of Vio molecules trapped in the binding sites of LHCII (see (Liu et al., 2004; Standfuss et al., 2005) for the structure). Such a process can be mediated, for example, via the light-driven trimer to monomer transformation of LHCII, reported to operate efficiently both in vitro and in chloroplasts (Garab et al., 2002). The light-driven molecular configuration changes of LHCII-bound Vio, associated with the “liberation” of xanthophyll molecules from the protein environment was demonstrated with the application of resonance Raman technique (Gruszecki et al., 2009; Grudzinski et al., 2016a). Interestingly, in contrast to Zea that adopts a roughly vertical, transmembrane orientation in the lipid bilayers (Grudzinski et al., 2017), the orientation of Vio is more heterogeneous and also comprises a pool of pigments oriented in the membrane plane (see Supplemental Figure 6). Such a heterogeneous distribution of Vio in the membranes, which can be related to the differences in number and localization of polar groups, seems to be particularly important from the standpoint of operation of the xanthophyll cycle. VDE belongs to a family of water-soluble lipocalin enzymes that after acidification of the environment dock in the lipid membrane, in the form of a dimer able to accommodate the entire xanthophyll molecule (Arnoux et al., 2009; Hallin et al., 2016). The mode of operation of the xanthophyll cycle and localization of VDE implies the necessity of Vio to adopt, at least temporarily, the planar orientation within the membrane (see Figure 7). On the other hand, molecules of newly synthesized Zea can readily adopt a roughly vertical orientation within the membrane, spanning the lipid bilayer (Supplemental Figure 6) (Grudzinski et al., 2017). Such a transmembrane localization and orientation can be further stabilized by specific or non-specific interactions of xanthophylls with interfacial regions of proteins. The importance of PsbS in stabilizing the transmembrane orientation of xanthophylls follows directly from the comparison of the results of the Raman photo-selection experiments carried out with the oriented thylakoid membranes isolated from the WT plants and *npq*4 mutants (Figure 1D). On the other hand, the results of the spectroscopic study presented here suggest an indirect role of PsbS in influencing xanthophyll localization and orientation, via controlling the supramolecular organization of the membrane proteins. Based on the results of the previous studies we can propose that structures of this type can be formed largely by LHCII (Zhou et al., 2020). A very important physiological aspect of the direct presence of Zea in the lipid phase of the thylakoid membranes is the protection against oxidative damage of polyunsaturated fatty acid chains (Havaux et al., 1991; Havaux and Niyogi, 1999).

**Figure 7.**
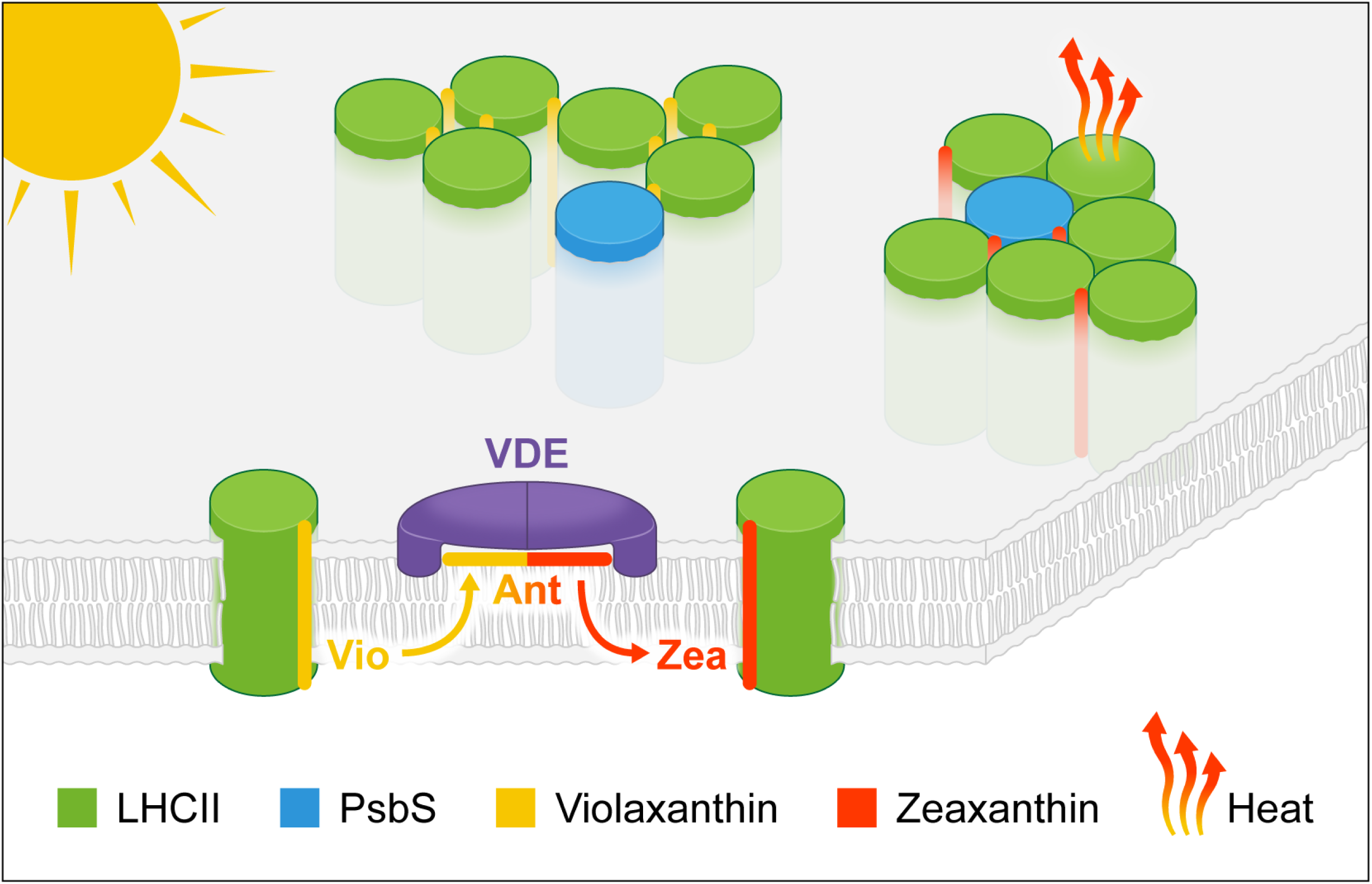
A model presenting localization of PsbS and the xanthophyll cycle pigments in the thylakoid membrane. The model presents also, discussed in the paper, the influence of PsbS and the xanthophyll cycle pigments on the molecular organization of selected pigment-protein complexes, important from the standpoint of photoprotection. VDE stands for violaxanthin de-epoxidase enzyme, Ant for antheraxanthin. See the text for a detailed description and discussion.

Importantly, the joint presence of Zea and PsbS in the thylakoid membranes has been demonstrated to be necessary to effectively develop the mechanism of photoprotective excitation quenching (Niyogi et al., 1998; Li et al., 2000; Sylak-Glassman et al., 2014) (Figure 1B, Figure 2C, Figure 3 and Figure 1C). The results of the fluorescence lifetime analysis of Chl *a* show that the thylakoid membranes lacking one of elements of such a dyad, either Zea or PsbS, are not able to effectively modulate the excitation quenching in response to changing light intensity (Figure 1C) (Sylak-Glassman et al., 2014). As briefly outlined in the introduction, one of the popular concepts regarding the photoprotective significance of the xanthophyll cycle is based on the mechanism of the direct quenching of excessive excitation energy by Zea. In the present work, we do not find arguments to support the mechanism that involves a direct Chl *a* excitation quenching via energy transfer to Zea. A similar conclusion has been drawn in some other recent studies, e.g. in the case of energy dissipation in intact *A. thaliana* leaves (Kress and Jahns, 2017) and photophysical examination of LHCII containing either Vio or Zea (Son et al., 2020a). On the contrary, a significant decrease in NPQ observed in the *npq*4 leaves (Figure 1B) and no light-dependent excitation quenching in the thylakoid membranes isolated from *npq*4 plants (Figure 1C) was observed despite the elevated accumulation of Zea (Figure 1A). This suggests the operation of a mechanism in which modulation of excitation quenching is rather realized in the thylakoid membranes indirectly, e.g. via the influence on a molecular organization of the pigment-protein complexes or by modifying their immediate environment (Ruban et al., 1997; Gruszecki et al., 2006; Johnson et al., 2011; Xu et al., 2015; Welc et al., 2016; Son et al., 2020b; Zhou et al., 2020). The results of the experiments have been presented, showing that both Zea (Ruban et al., 1997; Gruszecki et al., 2006; Zhou et al., 2020) and PsbS (Sacharz et al., 2017) influence molecular organization of LHCII. According to the concept of Sacharz et al., PsbS acts as “seeding” centres for an aggregate formation of LHCII (Sacharz et al., 2017). Our current interpretation of the results is consistent with the observation that Zea does not exchange for Vio in the internal binding sites of the antenna complexes but is located at the periphery of the pigment-proteins, helping to create excitation quenching centers (Xu et al., 2015). Recently, it was demonstrated that both Vio and Zea promote the formation of supramolecular structures of LHCII in the model lipid-protein membranes, influencing critically the Chl *a* excitation quenching capability (Zhou et al., 2020). Importantly, the efficiency of excitation quenching in the LHCII assemblies was lower in the presence of Vio, as compared to pure LHCII, but significantly enhanced in the structures formed in the presence of Zea (Zhou et al., 2020). Remarkably, the results of the Chl *a* fluorescence lifetime analyses show that the presence of PsbS also guarantees that at low light intensities the antenna complexes do not form aggregated structures, potentially leading to uncontrolled excitation quenching (Figure 1C, Figure 3). The fact that PsbS modulates molecular organization of the thylakoid membrane proteins also when the xanthophyll cycle pool is dominated by Vio (Figure 5) suggests a possible effect of PsbS in stabilizing the transmembrane orientation also by Vio present directly in the lipid phase of the thylakoid membranes. Importantly, the results of the precise analyses of the molecular organization of the Photosystem II antenna complexes in the thylakoid membranes carried out with the application of freeze-fracture electron microscopy, show the xanthophyll cycle-dependent reorganization of LHCII (Johnson et al., 2011). It has been demonstrated that LHCII is involved in the formation of large clusters, relatively loosely packed, when containing Vio and isolated from the dark-adapted leaves, or relatively densely packed, when containing Zea and isolated from the strong light-exposed leaves (Johnson et al., 2011). Such results are consistent with the concept regarding the separation of trimeric LHCII by Vio under low light conditions, to prevent the uncontrolled formation of aggregated structures that may be potentially involved in excitation quenching (Janik et al., 2016; Welc et al., 2016; Zhou et al., 2020) and with the conclusion that Zea can be involved in the formation of a certain type of LHCII aggregated structures characterized by effective excitation quenching (Ruban et al., 1997; Gruszecki et al., 2006; Janik et al., 2016; Welc et al., 2016; Zhou et al., 2020). Interestingly, the average Chl *a* fluorescence lifetimes in the thylakoid membranes isolated from leaves exposed to low light are at a level of ∼1 ns (Figure 1C, Figure 3) are substantially lower than in the case of the model LHCII-lipid membranes (2.14 ns, (Zhou et al., 2020)). It is, therefore, possible, that the effect of Vio on the molecular organization of LHCII, directly associated with the potential to quench excitations, is more complex and depends on a presence of other membrane constituents, very likely another proteins, possibly PsbS. On the other hand, the lowest average fluorescence lifetime of Chl *a*, determined in intact leaves (Sylak-Glassman et al., 2014) (see also Figure 2C) and the thylakoid membranes isolated from leaves exposed to high light (Figure 3) are at the level of ∼0.5 ns, very close to that one found in the model lipid membranes containing LHCII and Zea (0.49 ns) (Zhou et al., 2020). This shows that, most probably, the natural localization and orientation of Zea in the lipid membranes promotes the formation of supramolecular structures of LHCII, characterized by a pronounced excitation quenching. Importantly, the Chl *a* fluorescence lifetimes in LHCII in the quenched state in intact leaves (Sylak-Glassman et al., 2014), model lipid membranes (Zhou et al., 2020) and isolated thylakoid membranes (Figure 2) are at the level of 0.5 ns, substantially higher than in the case of the aggregated structures of LHCII formed in the detergent-free environment (∼0.2 ns) (Adams et al., 2018). This is a clear indication that simple aggregation of LHCII is rather not a mechanism involved in the photoprotective excitation quenching in the photosynthetic apparatus of plants. On the other hand, the results of the present study show that interplay of numerous factors, including the lipid membrane environment, the presence of PsbS and the presence of either Vio or Zea can modulate the immediate LHCII environment as well as formation of supramolecular structures of the protein, controlling critically the Chl *a* excitation quenching capability. The structures stabilized by Vio are characterized by a minimal excitation quenching, in contrast to the structures stabilized by Zea, which are characterized by highly efficient excitation quenching (see the model presented in Figure 7). In our opinion, it is highly probable that such a control of excitation quenching in LHCII is mechanistically realized via influencing the conformation and dynamics in the complex, directly associated with the photophysical processes, including excitation energy transfer to the S1 state of carotenoids (Son et al., 2020b; Son et al., 2020a). Importantly, the results of the experiments reported in the present work show that the activity of the xanthophyll cycle pigments in the thylakoid membranes is potentiated by PsbS promoting the transmembrane localization and orientation of both Vio and Zea, via influencing the molecular organization of the membrane proteins. The results of the CD analyzes reveal that PsbS interferes with the formation of the ordered supramolecular structures of pigment-protein complexes, giving rise to a strong excitonic and/or psi type signal. (Figure 6, Supplemental Figure 8). It is therefore highly probable that contribution of PsbS to the photoprotective activity is associated with the effect on molecular organization of pigment-protein complexes, resulting in irregular and less tight protein packing, thus providing a room for accommodation of the xanthophyll cycle pigments. In accordance with this interpretation, the lack of PsbS in the *npq*4 plants results in the increase of availability of Vio for enzymatic deepoxidation (Figure 1A). The ability to be incorporated into supramolecular structures of the proteins seems to be particularly important in the case of Zea shown to determine critically excitation quenching capability of Chl *a* in pigment-protein complexes. Therefore, it can be concluded that the cooperative influence of PsbS and Zea on the molecular organization of pigment-protein complexes determines the effective photoprotection in the photosynthetic apparatus of plants.

## Methods

### Plant material and growth conditions

*Arabidopsis thaliana* (L.), wild-type Columbia (Col-0), *npq1-2* mutant (N3771) and *npq4-1* mutant (N66021) seeds were obtained from the Nottingham Arabidopsis Stock Centre (NASC). Plants used in the experiments were grown in plastic pots (8 x 8 x 8 cm) filled with universal commercial soil mixture (COMPOSANA^®^, Compo, Poland). 60% field water capacity (FWC) was maintained during the experiment by daily water supplementation to maintain the mass of the pots. Plants were grown in a growth chamber under controlled environmental conditions: white light photon flux density of 150 μmol photons m^-2^s^-1^, photoperiod 8 h day and 16 h night, temperature 18 °C (night) and 22 °C (day). Air humidity was 60 %. The experiments were carried out on youngest, fully expanded leaves of 5 to 6 weeks-old plants. The efficiency of photochemical reactions in Photosystem II (Y(II)) of plants subjected to experiments is shown in Figure 2A.

### Pigment analysis

In order to monitor the operation of the xanthophyll cycle, leaves were illuminated for 30 min with white light from the white light-emitting LED lamp (Jansjo, IKEA) at selected intensities controlled by a distance from the light source and monitored by a portable photometer (Optel, Poland). Before illumination plants were dark-adapted for 30 min. For each analysis corresponding to the same light intensity, a leaf was detached from a different plant. Immediately after illumination, leaves were frozen via immersion in liquid nitrogen. Pigments were extracted from the frozen material with an acetonitrile : methanol (72:8, v:v) solvent mixture and then pigment extracts were filtered and subjected to HPLC analysis with application of a phase-reversed, C18-coated column (length 250 mm, internal diameter 4.6 mm, grain 5 µm). The HPLC mobile phase was: acetonitrile : methanol : water (72:8:3 v/v, solvent A) and acetonitrile : dichloromethane : methanol (54:28:18 v/v, solvent B). A linear gradient procedure between the solvents A and B, at a flow rate of 1.5 mL/min, was applied. Pigments were quantified based on the integration of HPLC chromatograms. The system was calibrated with equimolar mixture of the photosynthetic pigment standards.

### Determination of a non-photochemical quenching

Determination of a non-photochemical quenching parameter (NPQ) and the efficiency of PSII photochemistry (Y(II)) were based on measurements with the application of the system consisting of GFS3000, Dual PAM 100 and 3010-Dual cuvette (Walz GmbH, Germany). The system uses a combination of the blue-light-emitting and the red-light-emitting diodes as an actinic light. Parameters were calculated as described elsewhere (Maxwell and Johnson, 2000). During the measurements of chlorophyll fluorescence in the leaf chamber of the gas exchange system leaf temperature was maintained within the range 22 ± 0.2 °C (irrespectively on light intensity), relative air humidity and CO_2_ concentration were 60% and 400 ppm. The actinic light source was a mixture of red and blue LEDs (75:25). The spectra of light sources are shown in Supplemental Figure 9. Each of NPQ values was determined in leaves after 30 min of dark adaptation and following illumination with actinic light of specified intensity for 30 min (n=4).

### Thylakoid membrane isolation and preparation of multilayers

Thylakoid membranes were isolated from leaves of *Arabidopsis thaliana* (wild type, *npq*1 and *npq*4). Leaves were illuminated with light intensity in the range 100-700 μmol photons m^-2^ s^-1^ for 30 min. Before illumination plants were stored in darkness for 30 min. Directly after illumination, leaves were ground in ice-cold buffer containing 20 mM Tricine (pH 7.6), 0.4 M sorbitol, filtered and centrifuged for 5 min at 4°C and 5000 *g.* The pellet was resuspended in 20 mM Tricine (pH 7.6). Ordered thylakoid multilayers were formed on a solid support (either quartz glass in the case of Raman or diamond in the case of FTIR measurements) via evaporation of the membrane suspension under argon atmosphere, according to the procedure elaborated, tested and described previously (Janik et al., 2013).

### Fluorescence Lifetime Imaging Microscopy

Fluorescence Lifetime Imaging Microscopy (FLIM) analyses were performed using confocal MicroTime 200 (PicoQuant, Germany) system coupled to an inverted microscope OLYMPUS IX71. The samples were excited with a 470 nm pulse laser, with a repetition rate 20 MHz and observation was made through 690/70 bandpass filter. In order to prepare samples containing oriented multi-bilayer membranes, thylakoids were deposited on non-fluorescent glass slides (Menzel-Glaser) and partially dried in an argon atmosphere. The samples were excited with a 470 nm pulse laser, with a repetition rate 20 MHz. The lifetime resolution was better than 16 ps. Fluorescence photons were collected with a 100x oil objective (NA 1.3, OLYMPUS UPlanSApo) and confocal pinhole of 50 μm in diameter was used. The scattered light was removed using ZT473RDC XT dichroic filter (Analysentechnik), 470 notch filter (Semrock) and observation was made through 690/70 bandpass filter. Results of measurements were analyzed using SymPhoTime 64 software.

### Fluorescence Lifetime Measurements

Time-resolved fluorescence intensity decays were measured using FluoTime 300 spectrometer (PicoQuant, Germany) in the so-called front face configuration providing measurements for whole leaves pre-treated with appropriate illumination. The light generated from pulsed solid state laser operating at 635 nm and frequency of 20 MHz was used to excite the sample. The emission was filtered by 665 glass, long wavelength pass filter (Edmund Optics), 680/13 band pass filter (Semrock), monochromator set at 680 nm and polarizer at the magic angle to excitation. The detection was accomplished by a microchannel plate in time-correlated single photon counting mode. Fluorescence lifetime decays were fitted using FluoFit Pro software (PicoQuant, Germany).

Intact leaves were detached from plants, placed on moist filter paper and dark adapted for 30 min. Directly before fluorescence lifetime measurements, leaves were illuminated with a white light LED lamp with the controlled photon flux density. A measurement of a single leaf lasted typically between 1 and 2 min.

### Raman spectroscopy

Raman spectra recording and imaging were carried out using an inVia Reflex confocal Raman microscope (Renishaw, UK) with an argon laser (Stellar-REN, Modu-Laser™, USA) operating either at 457 nm (0.35 mW laser power at the sample) or at 514 nm (1.2 mW). We used 20x long working distance objective (SLMPlan N, NA=0.25, Olympus) both for imaging of the thylakoid membranes and intact leaves. In the case of intact leaves, they were detached from plants, placed at moisture filter paper and dark-adapted for 30 min. Directly before Raman imaging leaves were illuminated (at room temperature), with the white light LED lamp at controlled photon flux density. Owing to the fact that imaging of a single sample lasted for ca. 19 min, in order to stop light-driven enzymatic reactions, samples were cooled down to -30 °C and this temperature were stabilized during the measurements. In the case of thylakoid multilayers membranes, the samples were analyzed at room temperature. The other parameters were the same as used for intact leaves. The imaged area was 260 µm x 400 µm with single pixel size 4 µm x 4 µm. At each point, the spectra were recorded at about 1 cm^-1^ spectral resolution (2400 lines/mm grating) in the spectral region 350÷1950 cm^-1^ using EMCCD Newton 970 detection camera from Andor, UK. The detector was cooled to -50 °C. Images were acquired by use of the Renishaw WiRE 4.4 system in high-resolution mapping mode (HR maps). The acquisition time for a single spectrum in each point was 0.05 s. All spectra were pre-processed by cosmic ray removing, noise filtering and baseline correction using WiRE 4.4 software from Renishaw, UK. The spectra corresponding to single chloroplasts were extracted from the map and subjected to analysis.

### Fourier transform infrared spectroscopy

Fourier transform infrared (FTIR) spectra were recorded with Nicolet iS50 FTIR spectrometer (Thermo Scientific, USA), equipped with a single reflection diamond ATR cell. Thylakoid samples were deposited at the diamond surface by evaporation under an argon stream. Absorption spectra were collected in the region between 4000 and 600 cm^−1^ with a 4 cm^-1^ resolution.

### Circular Dichroism spectroscopy

Circular Dichroism (CD) measurements were performed using Chirascan-plus spectrometer (Applied Photophysics, UK) in the range of 300–750 nm, at room temperature. The spectra were collected from the thylakoids suspended in buffer (20mM Tricine, 10mM KCl, pH=7.6). All the samples were diluted before measurements to obtain the identical absorbance of 0.1 at the absorption maximum in the red spectral region.

## Supplemental Data

**Supplemental Figure 1.** A scheme of the reactions of the xanthophyll cycle in higher plants.

**Supplemental Figure 2.** Light intensity dependence of zeaxanthin level in leaves of *Delonix regia*.

**Supplemental Figure 3.** Light intensity dependence of zeaxanthin accumulation in plants.

**Supplemental Figure 4.** Raman spectroscopy analysis of multilayers formed with the thylakoid membranes.

**Supplemental Figure 5.** A model of the carotenoid-containing lipid bilayer subjected to a Raman scattering analysis.

**Supplemental Figure 6.** Fluorescence microscopy images of single lipid vesicles containing xanthophylls.

**Supplemental Figure 7.** FTIR spectra of the thylakoid membranes.

**Supplemental Figure 8.** CD spectra of the thylakoid membranes.

**Supplemental Figure 9.** Spectra of the lamps used as actinic light sources.

**Supplemental Table 1.** Xanthophyll composition in leaves.

## ACKNOWLEDGEMENTS

National Science Center, Poland is acknowledged for financial support within the project 2016/22/A/NZ1/00188. The research was carried out with the equipment purchased thanks to the financial support of the European Regional Development Fund in the framework of the Development of Eastern Poland Operational Program.

## AUTHOR CONTRIBUTIONS

W.I.G. conceived the project with R.L., W.G. and R.W., R.W., R.L., D.K., M.Z., M.M., E.R., K.S., R.M., A.N. performed the experiments. All the authors analyzed the data. W.I.G. wrote the manuscript with input from other coauthors.

## Supplemental Information

**Supplemental Figure 1.**
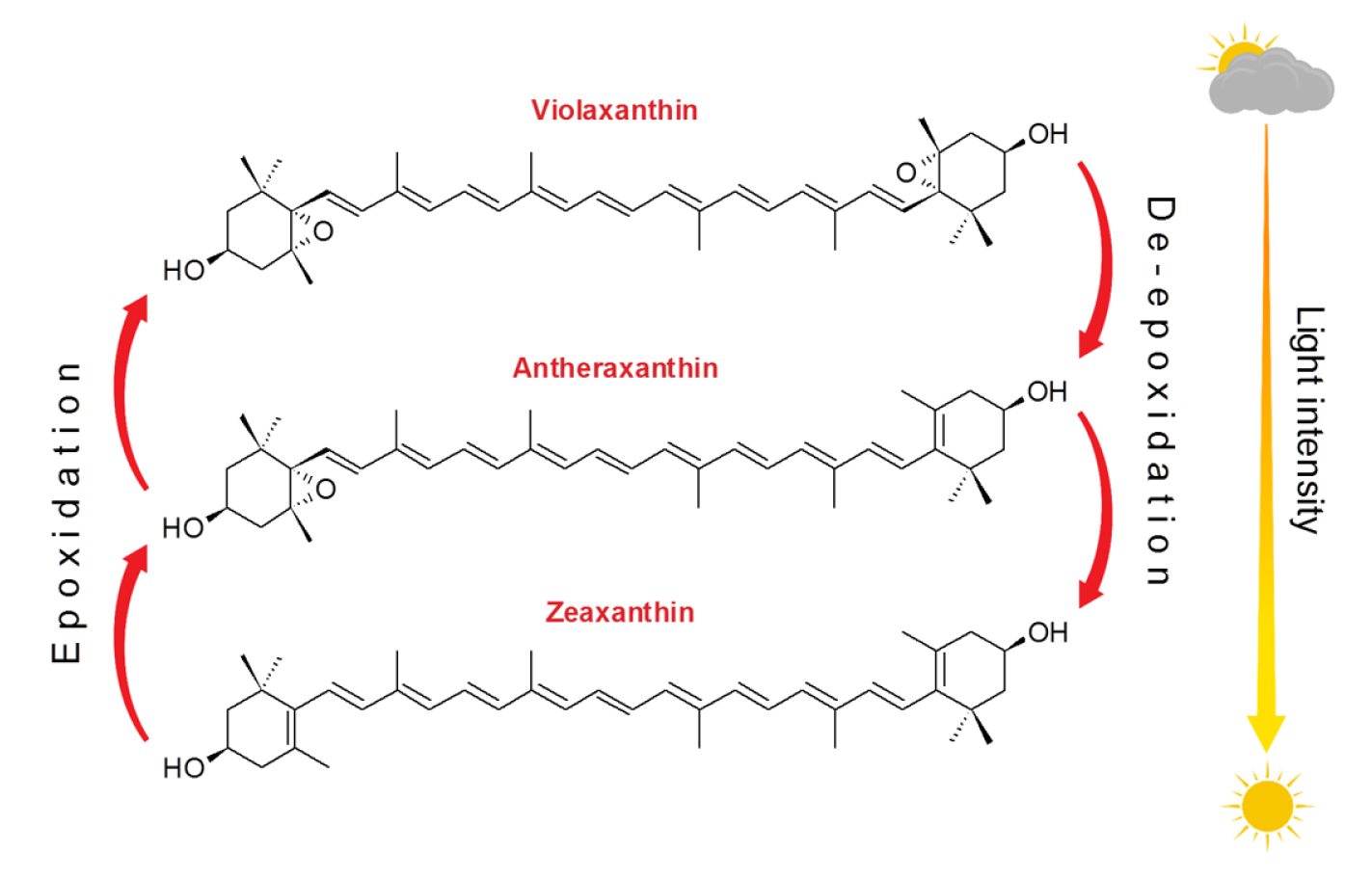
Scheme of the xanthophyll cycle in higher plants.

**Supplemental Figure 2.**
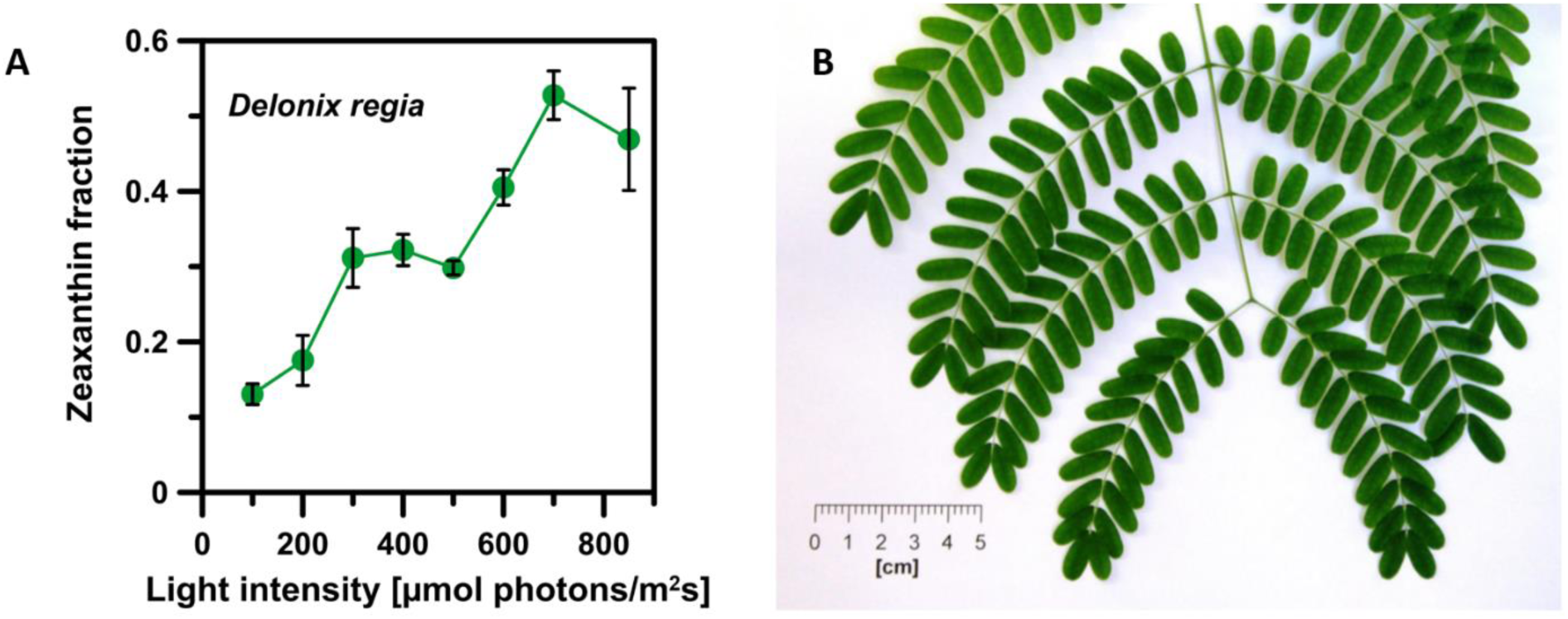
Light intensity dependence of zeaxanthin level in leaves of *Delonix regia*. (A) Zeaxanthin level in leaves subjected to illumination for 30 min with white light at different intensities (represented in the abscissa axis). Zeaxanthin level was calculated as a fraction of the xanthophyll cycle pigments Zea/(Vio+Ant+Zea). The results represent mean from 6-9 experiments ± S.D. (B) Photo of *Delonix regia*. *Delonix regia* plants were cultivated under room conditions with natural light. The majority of measurements were carried out during springtime. The photoperiod was approximately 13 h (day) and 11 h (night). The room temperature and photon flux density were in the range of 21-23 °C and 100-150 μmol photons m^-2^ s^-1^ at noon. *Delonix regia* has been selected to be compared to *Arabidopsis thaliana* as an essentially different plant model: a tropical tree native to Africa (*D*. *regia*) versus a small flowering plant native to Eurasia (*A. thaliana*).

**Supplemental Figure 3.**
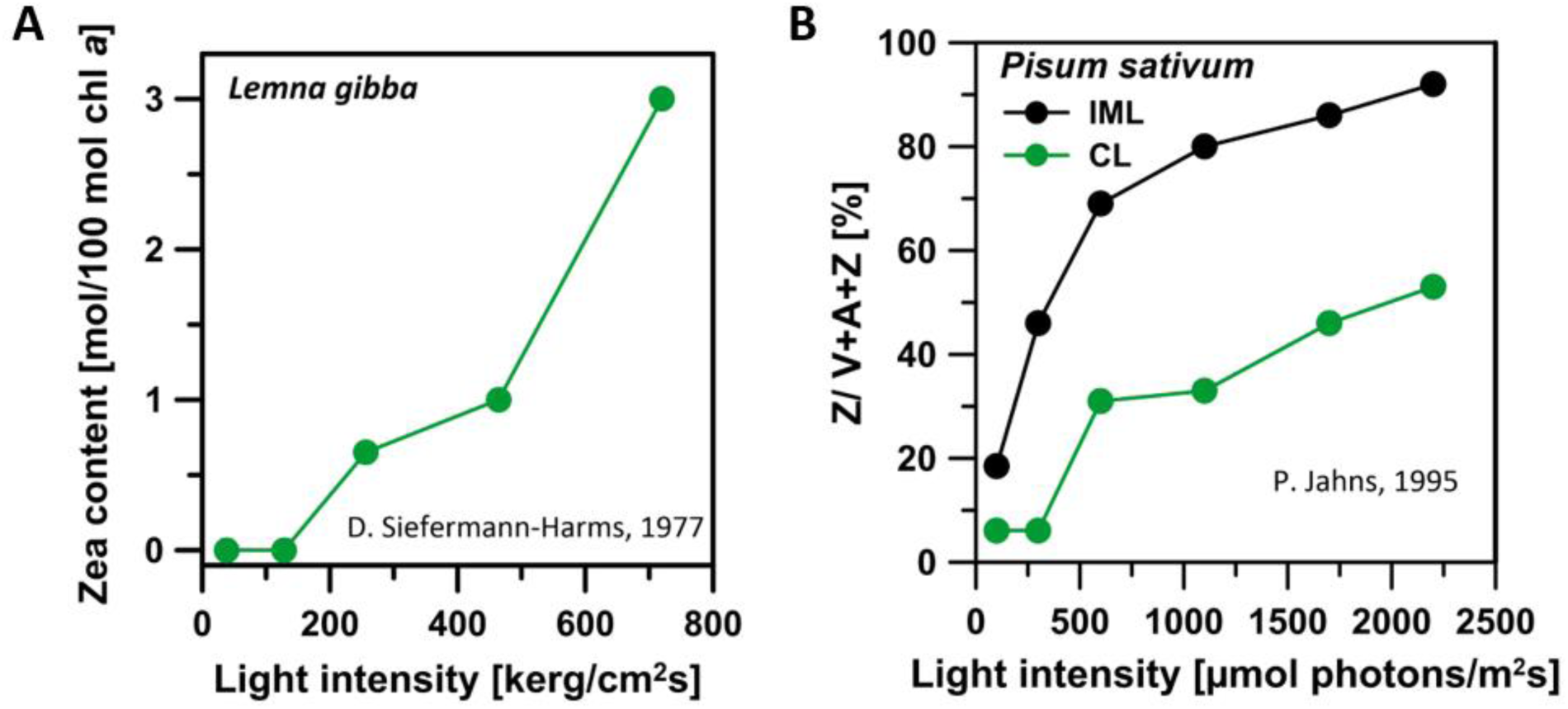
Light intensity dependence of zeaxanthin accumulation in plants. **(A)** The dependency drawn based on the original results published by D. Siefermann-Harms, The xanthophyll cycle in higher plants, in: M. Tevini, H.K. Lichtenthaller (Eds.) Lipids and Lipid Polymers in Higher Plants, Springer-Verlag, Berlin, 1977, pp. 218-230 (Figure 10a). Light intensity units are used as in the original publication. (B) The dependencies are drawn based on the original results published by P. Jahns, The Xanthophyll Cycle in Intermittent Light-Grown Pea-Plants - Possible Functions of Chlorophyll a/b Binding-Proteins, Plant Physiology, 108 (1995) 149-156 (Figure 1 and Figure 2). Two types of plants were examined: grown under continuous light (CL) and intermittent light (IML) conditions.

**Supplemental Figure 4.**
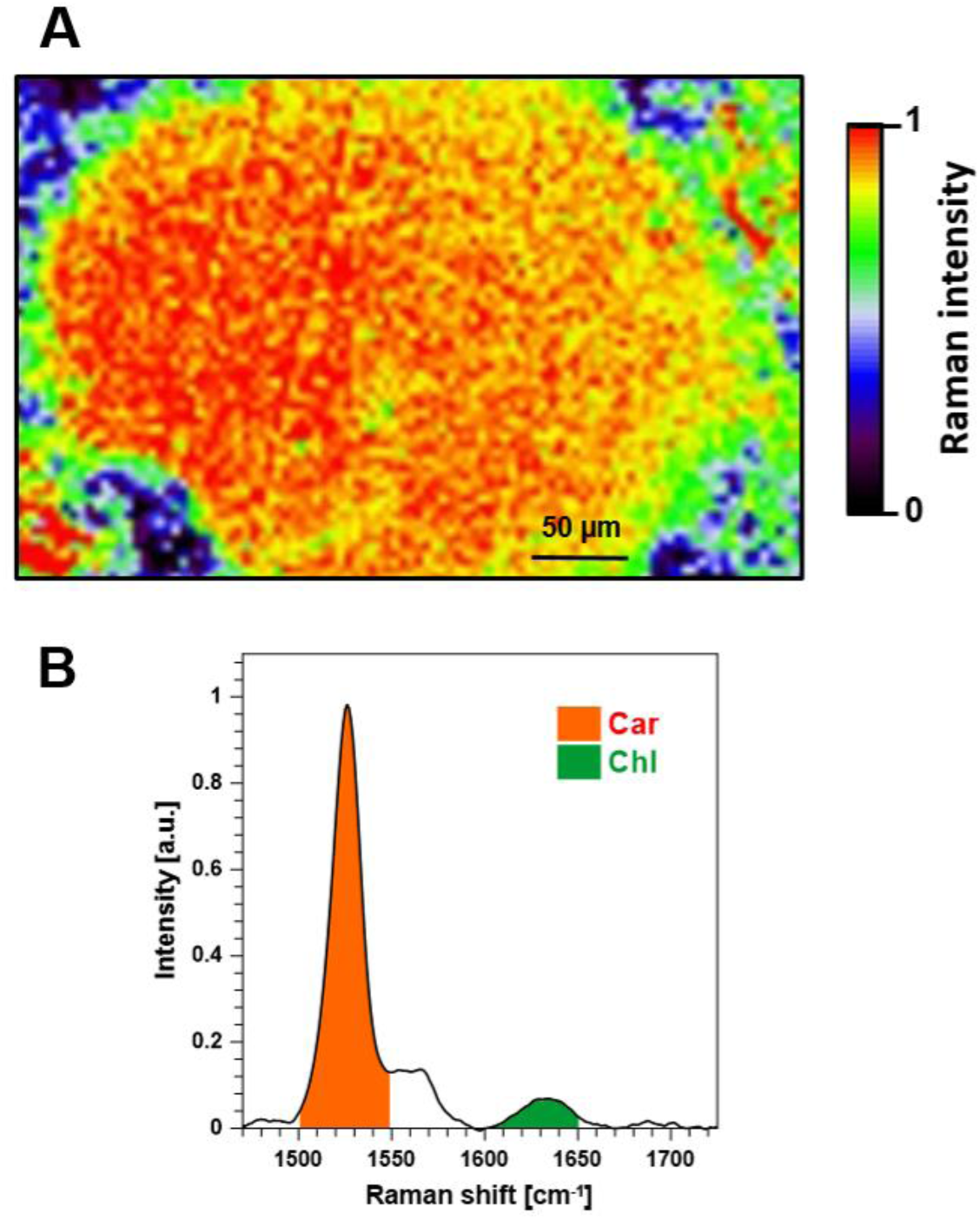
Raman spectroscopy analysis of multilayers formed with the thylakoid membranes. Raman analysis was performed with the application of the 457 nm laser line being in resonance with both carotenoids and chlorophylls. Panel A shows an exemplary Raman image of the multilayer formed with the thylakoid membranes isolated from *A. thaliana* WT, pre-illuminated for 30 min with white light, 700 µmol photons/m^2^s. Color codes represent Raman intensity determined as the integration of the spectra in the region corresponding to the C=C stretching vibrations of carotenoids (1500-1550 cm^-1^). The imaged area was 260 µm x 400 µm with single pixel size 4 µm x 4 µm. The exemplary spectrum recorded in a single pixel of the sample is presented in panel B. The spectral components representing the C=C stretching of carotenoids (1500-1550 cm^-1^) and chlorophylls (1600-1650 cm^-1^) are marked by shaded areas beneath the spectrum. The ratio Car/Chl values, representing the contribution of carotenoids relative to chlorophylls in the resonance Raman spectra of leaves, were calculated by dividing the integration results of the bands assigned to Car and Chl.

**Supplemental Figure 5.**
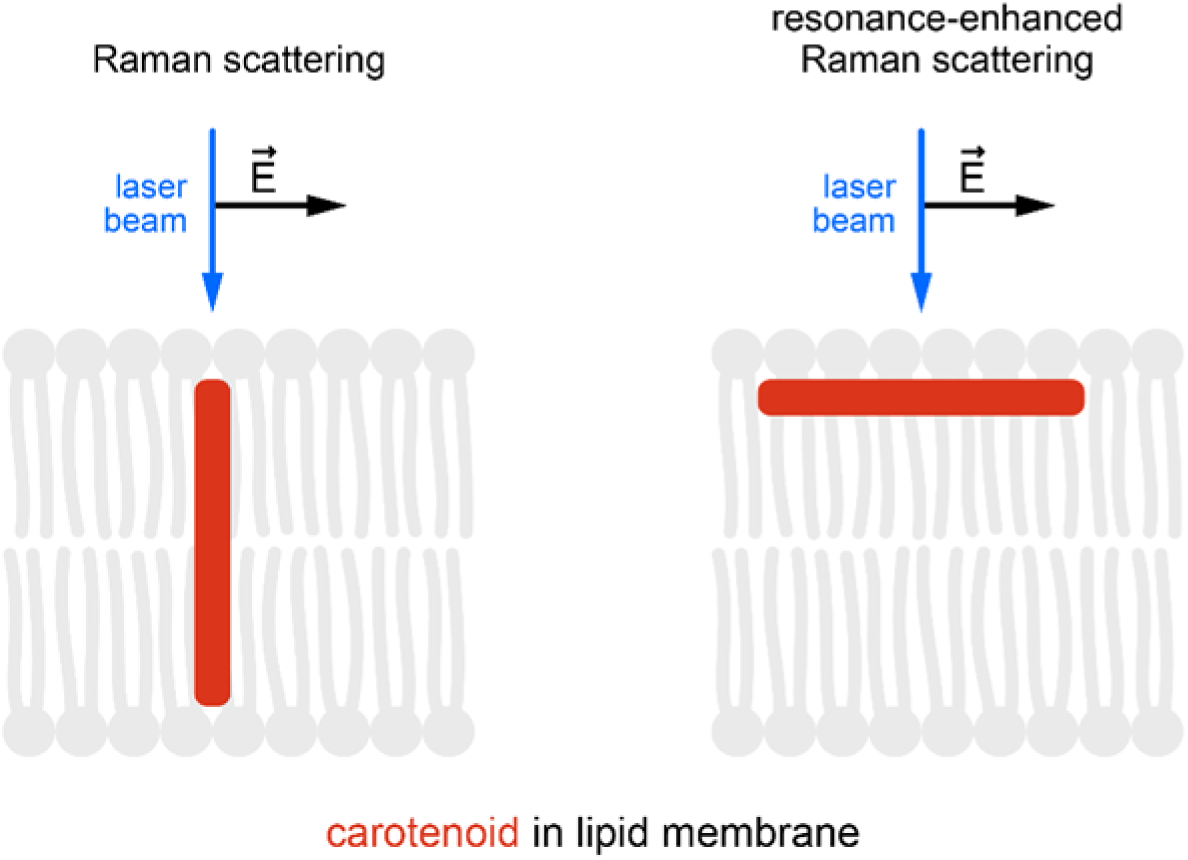
A model of the carotenoid-containing lipid bilayer subjected to a Raman scattering analysis. The model shows two orientations of the carotenoid chromophore with respect to the membrane plane, critically different from the standpoint of the intensity of Raman scattering. The chromophore oriented along the direction of the laser beam gives rise to Raman scattering signal, in contrast to the pool oriented in the membrane plane, which can give rise to a significantly enhanced resonance Raman signal.

**Supplemental Figure 6.**
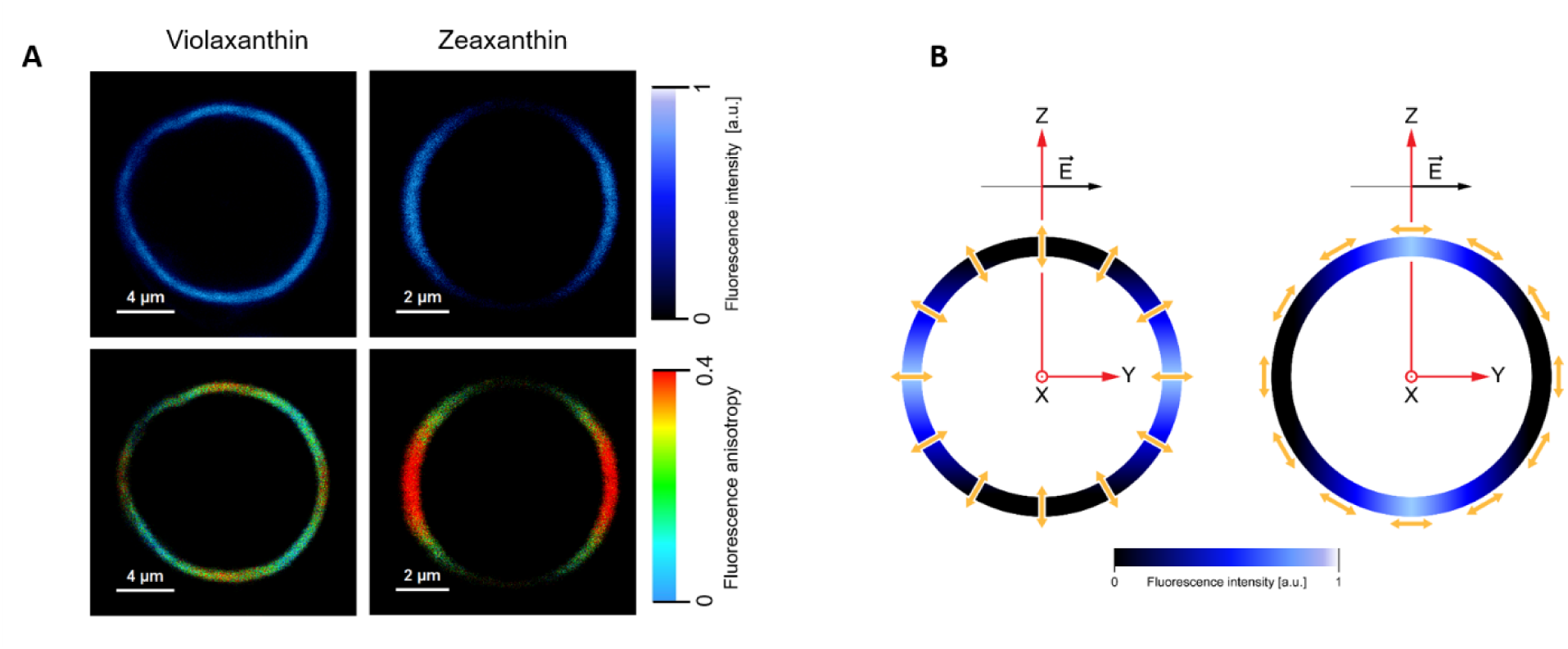
Fluorescence microscopy images of single lipid vesicles containing xanthophylls. Each image, generated with a confocal microscopy technique, presents an equatorial cross-section of a vesicle. Imaging is based upon fluorescence of carotenoids. (A) Upper panels show images of single giant unilamellar vesicles (GUV) formed with dipalmitoyl phosphatidylcholine (DPPC) containing 0.5 mol% of violaxanthin or zeaxanthin (indicated). Lower panels show fluorescence anisotropy images of the same structures. (B) The scheme presenting an idea of photoselection. Molecules oriented along the axis normal to the plane of the membrane give rise to strong fluorescence signal and high fluorescence anisotropy in the membrane regions located in the left-hand side and the right-hand side of the vesicle. Molecules oriented in the membrane plane give rise to strong fluorescence and high fluorescence anisotropy in the membrane regions located in the upper and lower regions of the vesicle. The images presented are consistent with the picture regarding the molecular distribution of xanthophylls according to which Zea adopts preferentially a vertical orientation within the lipid bilayer in contrast to Vio that can adopt also a horizontal orientation. A methodological background of this experimental approach and data interpretation was described previously (Grudzinski, W., Nierzwicki, L., Welc, R., Reszczynska, E., Luchowski, R., Czub, J., and Gruszecki, W.I. (2017). Localization and Orientation of Xanthophylls in a Lipid Bilayer. Sci. Rep. 7, 9619.).

**Supplemental Figure 7.**
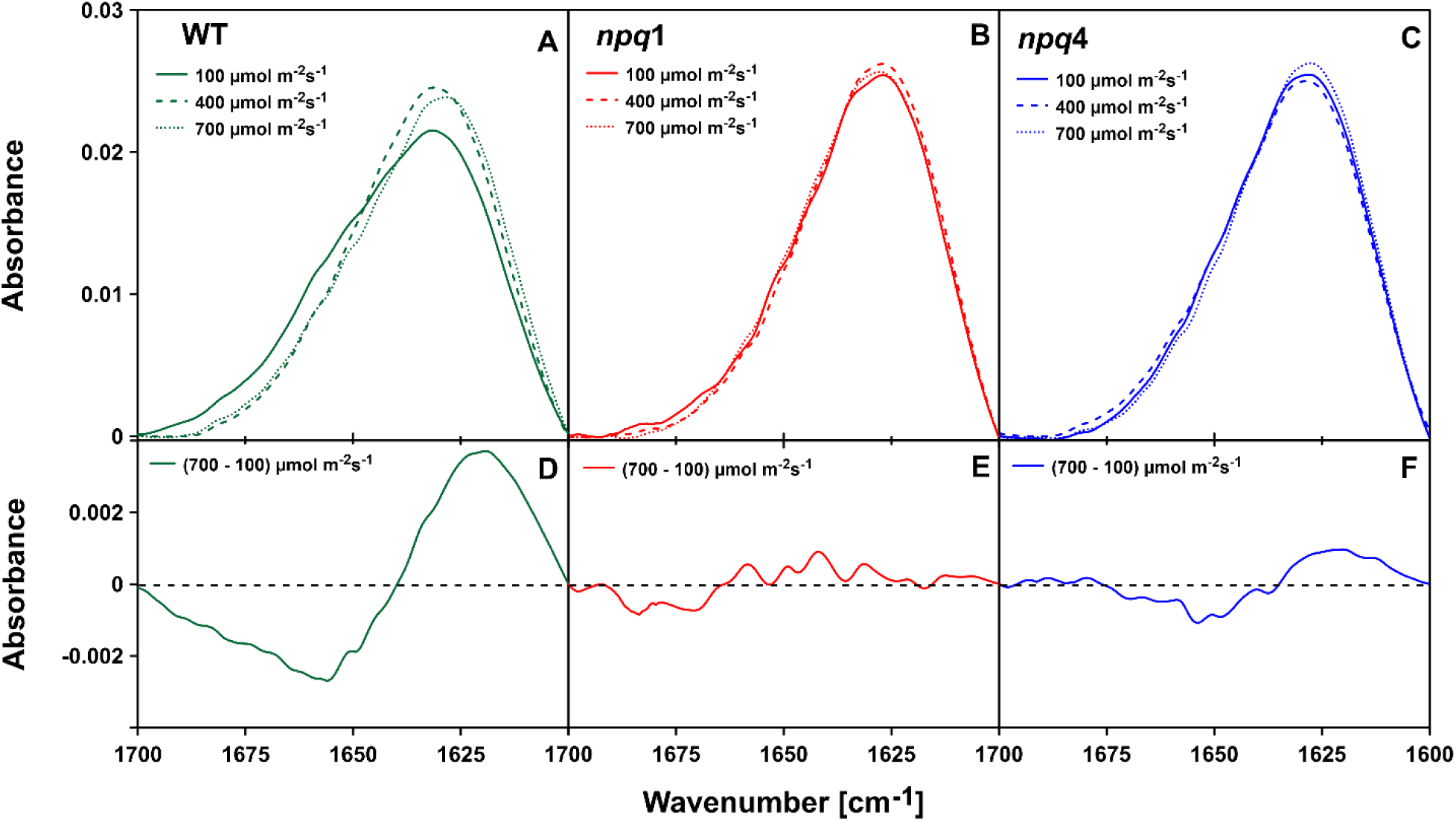
FTIR spectra of the thylakoid membranes. The infrared absorption spectra were recorded from the thylakoid membranes of three genotypes of *A. thaliana*: WT, *npq*1 and *npq*4, isolated from leaves pre-illuminated for 30 min with a white light of different intensities (indicated). Panels A, B and C present the original spectra and the panels D, E and F present the difference spectra. The original, surface-normalized spectra are presented in the amide I region (1600-1700 cm^-1^). The samples were dark-adapted under argon for 15 min before and during the measurements. Each spectrum in panels A, B and C represents a different experiment.

**Supplemental Figure 8.**
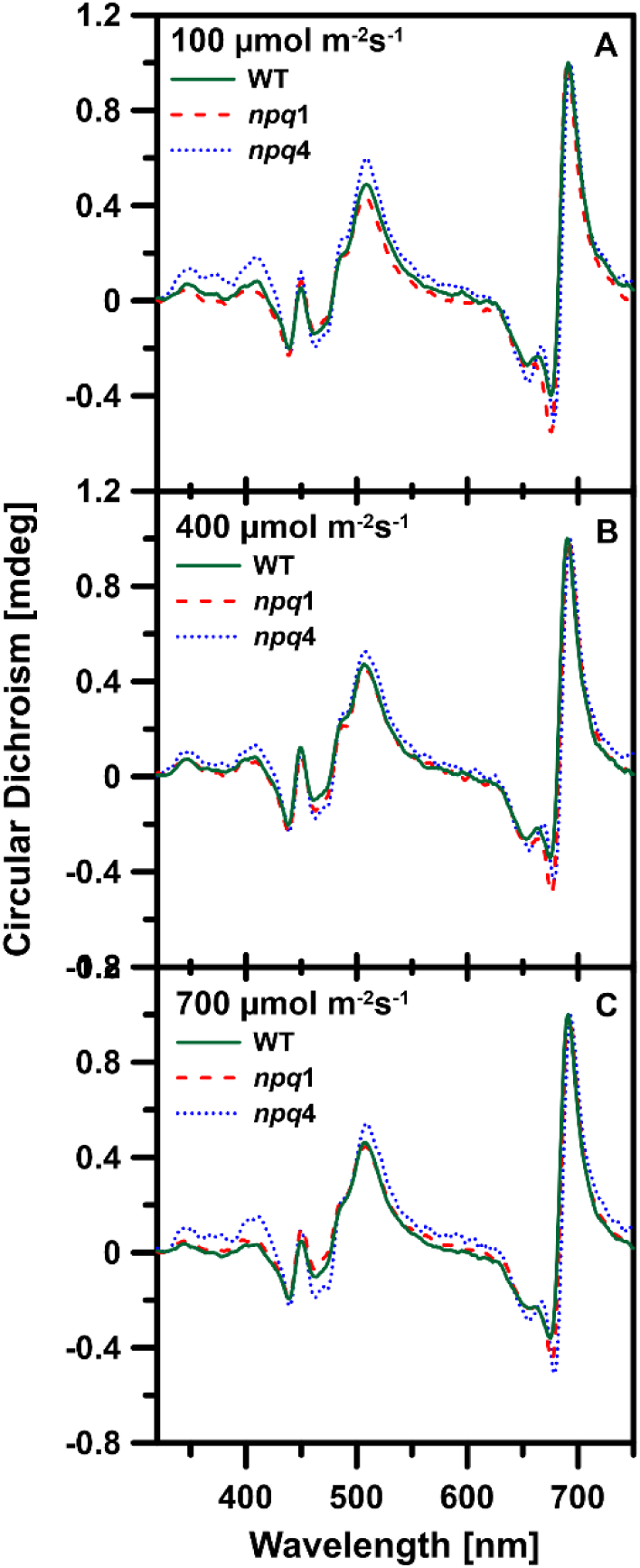
CD spectra of the thylakoid membranes. The circular dichroism spectra were recorded from the thylakoid membranes of three genotypes of *A. thaliana*: WT, *npq*1 and *npq*4, isolated from leaves pre-illuminated for 30 min with a white light of different intensities (indicated). The samples were dark-adapted for 15 min before the measurements. The spectra were normalized at the maximum. Each spectrum represents a different experiment.

**Supplemental Figure 9.**
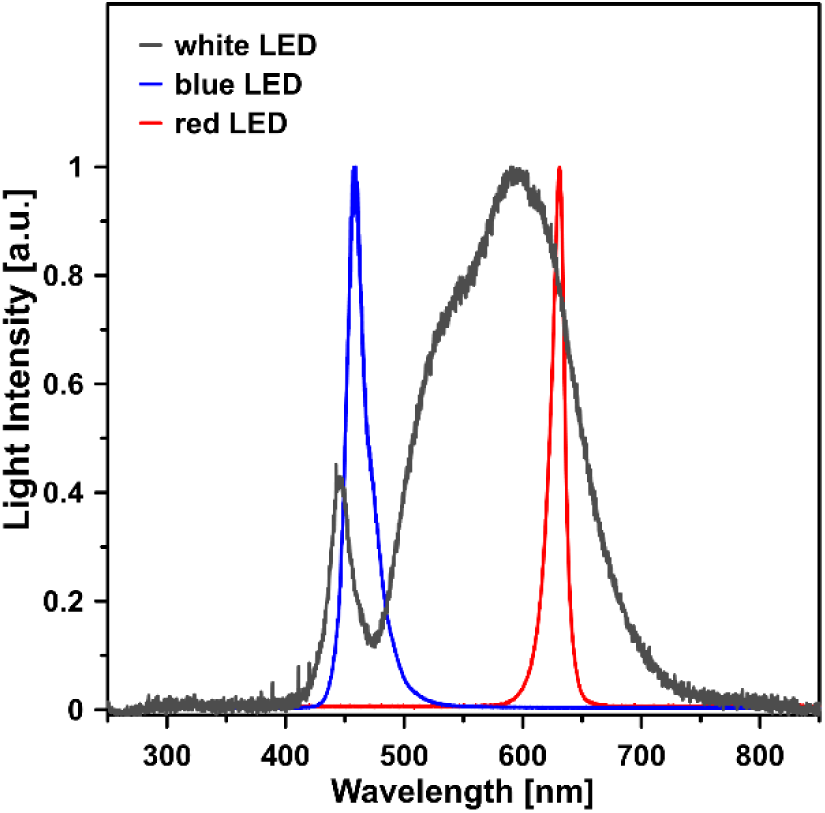
Spectra of the lamps used as actinic light sources. The spectra were normalized at the maximum. In all the experiments the white LED lamp was used as an actinic light source except for the data presented in Figure 1B and Figure 2A. In such a case, a mixture (75:25) of the red LED and blue LED actinic light sources was used.

Supplemental Table 1. Xanthophyll composition in leaves.

**Supplemental Table 1. Xanthophyll composition in leaves**
| Xanthophyll | *A. thaliana* WT | *A. thaliana npq1* | *A. thaliana npq4* |
| --- | --- | --- | --- |
| Neo | 37.87±1.67 | 39.75±2.77 | 40.18±2.73 |
| Lut | 117.68±6.38 | 121.76±3.72 | 127±6.24 |
| Vio+Ant+Zea | 25.79±3.38 | 34.58±3.33 | 24.78±3.94 |

Note: The xanthophyll content is given in relation to 1000 Chl (*a* + *b*). Neo stands for neoxanthin, Vio for violaxanthin, Ant for antheraxanthin, Zea for Zeaxanthin, Lut for lutein. Pigment level determined for dark-adapted leaves ± S.D. (n=4).

## REFERENCES

1. Adams, P.G., Vasilev, C., Hunter, C.N., and Johnson, M.P. (2018). Correlated fluorescence quenching and topographic mapping of Light-Harvesting Complex II within surface-assembled aggregates and lipid bilayers. Biochim. Biophys. Acta 1859, 1075–1085.

2. Arnoux, P., Morosinotto, T., Saga, G., Bassi, R., and Pignol, D. (2009). A structural basis for the pH-dependent xanthophyll cycle in Arabidopsis thaliana. Plant Cell 21, 2036–2044.

3. Bonente, G., Howes, B.D., Caffarri, S., Smulevich, G., and Bassi, R. (2008). Interactions between the photosystem II subunit PsbS and xanthophylls studied in vivo and in vitro. J. Biol. Chem. 283, 8434–8445.

4. Chmeliov, J., Gelzinis, A., Songaila, E., Augulis, R., Duffy, C.D.P., Ruban, A.V., and Valkunas, L. (2016). The nature of self-regulation in photosynthetic light-harvesting antenna. Nat Plants 2, 16045.

5. Croce, R., and van Amerongen, H. (2014). Natural strategies for photosynthetic light harvesting. Nature Chemical Biology 10, 492–501.

6. Fan, M.R., Li, M., Liu, Z.F., Cao, P., Pan, X.W., Zhang, H.M., Zhao, X.L., Zhang, J.P., and Chang, W.R. (2015). Crystal structures of the PsbS protein essential for photoprotection in plants. Nat. Struct. Mol. Biol. 22, 729–U115.

7. Frank, H.A., Bautista, J.A., Josue, S.J., and Young, A.J. (2000). Mechanism of nonphotochemical quenching in green plants: energies of the lowest excited singlet states of violaxanthin and zeaxanthin. Biochemistry 39, 2831–2837.

8. Garab, G., Cseh, Z., Kovacs, L., Rajagopal, S., Varkonyi, Z., Wentworth, M., Mustardy, L., Der, A., Ruban, A.V., Papp, E., Holzenburg, A., and Horton, P. (2002). Light-induced trimer to monomer transition in the main light-harvesting antenna complex of plants: thermo-optic mechanism. Biochemistry 41, 15121–15129.

9. Grudzinski, W., Matula, M., Sielewiesiuk, J., Kernen, P., Krupa, Z., and Gruszecki, W.I. (2001). Effect of 13-cis violaxanthin on organization of light harvesting complex II in monomolecular layers. Biochim. Biophys. Acta 1503, 291–302.

10. Grudzinski, W., Nierzwicki, L., Welc, R., Reszczynska, E., Luchowski, R., Czub, J., and Gruszecki, W.I. (2017). Localization and orientation of xanthophylls in a lipid bilayer. Sci. Rep. 7, 9619.

11. Grudzinski, W., Janik, E., Bednarska, J., Welc, R., Zubik, M., Sowinski, K., Luchowski, R., and Gruszecki, W.I. (2016a). Light-driven reconfiguration of a xanthophyll violaxanthin in the photosynthetic pigment-protein complex LHCII: A resonance Raman study. J. Phys. Chem. B 120, 4373–4382.

12. Grudzinski, W., Janik, E., Bednarska, J., Welc, R., Zubik, M., Sowinski, K., Luchowski, R., and Gruszecki, W.I. (2016b). Light-Driven Reconfiguration of a Xanthophyll Violaxanthin in the Photosynthetic Pigment-Protein Complex LHCII: A Resonance Raman Study. J Phys Chem B 120, 4373–4382.

13. Gruszecki, W.I., Grudzinski, W., Gospodarek, M., Patyra, M., and Maksymiec, W. (2006). Xanthophyll-induced aggregation of LHCII as a switch between light-harvesting and energy dissipation systems. Biochim. Biophys. Acta 1757, 1504–1511.

14. Gruszecki, W.I., Gospodarek, M., Grudzinski, W., Mazur, R., Gieczewska, K., and Garstka, M. (2009). Light-induced change of configuration of the LHCII-bound xanthophyll (tentatively assigned to violaxanthin): A resonance Raman study. J. Phys. Chem. B 113, 2506–2512.

15. Gruszecki, W.I., Stiel, H., Niedzwiedzki, D., Beck, M., Milanowska, J., Lokstein, H., and Leupold, D. (2005). Towards elucidating the energy of the first excited singlet state of xanthophyll cycle pigments by X-ray absorption spectroscopy. Biochim. Biophys. Acta 1708, 102–107.

16. Hallin, E.I., Hasan, M., Guo, K., and Akerlund, H.E. (2016). Molecular studies on structural changes and oligomerisation of violaxanthin de-epoxidase associated with the pH-dependent activation. Photosynth. Res. 129, 29–41.

17. Hartel, H., Lokstein, H., Grimm, B., and Rank, B. (1996). Kinetic studies on the xanthophyll cycle in barley leaves - Influence of antenna size and relations to nonphotochemical chlorophyll fluorescence quenching. Plant Physiol. 110, 471–482.

18. Havaux, M., and Niyogi, K.K. (1999). The violaxanthin cycle protects plants from photooxidative damage by more than one mechanism. Proc. Natl. Acad. Sci. USA 96, 8762–8767.

19. Havaux, M., Gruszecki, W.I., Dupont, I., and Leblanc, R.M. (1991). Increased heat emission and its relationship to the xanthophyll cycle in pea leaves exposed to strong light stress. J. Photochem. Photobiol. B: Biol. 8, 361–370.

20. Havaux, M., Eymery, F., Porfirova, S., Rey, P., and Dormann, P. (2005). Vitamin E protects against photoinhibition and photooxidative stress in Arabidopsis thaliana. Plant Cell 17, 3451–3469.

21. Holt, N.E., Zigmantas, D., Valkunas, L., Li, X.P., Niyogi, K.K., and Fleming, G.R. (2005). Carotenoid cation formation and the regulation of photosynthetic light harvesting. Science 307, 433–436.

22. Holzwarth, A.R., Miloslavina, Y., Nilkens, M., and Jahns, P. (2009). Identification of two quenching sites active in the regulation of photosynthetic light-harvesting studied by time-resolved fluorescence. Chem. Phys. Lett. 483, 262–267.

23. Jahns, P. (1995). The xanthophyll cycle in intermittent light-grown pea-plants - possible functions of chlorophyll a/b binding-proteins. Plant Physiol. 108, 149–156.

24. Jahns, P., and Holzwarth, A.R. (2012). The role of the xanthophyll cycle and of lutein in photoprotection of photosystem II. Bba-Bioenergetics 1817, 182–193.

25. Jahns, P., Latowski, D., and Strzalka, K. (2009). Mechanism and regulation of the violaxanthin cycle: The role of antenna proteins and membrane lipids. Biochim. Biophys. Acta 1787, 3–14.

26. Janik, E., Bednarska, J., Zubik, M., Sowinski, K., Luchowski, R., Grudzinski, W., Matosiuk, D., and Gruszecki, W.I. (2016). The xanthophyll cycle pigments, violaxanthin and zeaxanthin, modulate molecular organization of the photosynthetic antenna complex LHCII. Arch. Biochem. Biophys. 592, 1–9.

27. Janik, E., Bednarska, J., Zubik, M., Puzio, M., Luchowski, R., Grudzinski, W., Mazur, R., Garstka, M., Maksymiec, W., Kulik, A., Dietler, G., and Gruszecki, W.I. (2013). Molecular architecture of plant thylakoids under physiological and light stress conditions: A study of lipid-light-harvesting complex II model membranes. Plant Cell 25, 2155–2170.

28. Johnson, M.P., Goral, T.K., Duffy, C.D.P., Brain, A.P.R., Mullineaux, C.W., and Ruban, A.V. (2011). Photoprotective energy dissipation involves the reorganization of Photosystem II light-harvesting complexes in the grana membranes of spinach chloroplasts. Plant Cell 23, 1468–1479.

29. Krause, G.H., and Weis, E. (1984). Chlorophyll fluorescence as a tool in plant physiology .2. Interpretation of fluorescence signals. Photosynth. Res. 5, 139–157.

30. Kress, E., and Jahns, P. (2017). The dynamics of energy dissipation and xanthophyll conversion in Arabidopsis Indicate an indirect photoprotective role of zeaxanthin in slowly inducible and relaxing components of non-photochemical quenching of excitation energy. Front. Plant Sci. 8.

31. Latowski, D., Akerlund, H.E., and Strzalka, K. (2004). Violaxanthin de-epoxidase, the xanthophyll cycle enzyme, requires lipid inverted hexagonal structures for its activity. Biochemistry 43, 4417–4420.

32. Li, X.P., Bjorkman, O., Shih, C., Grossman, A.R., Rosenquist, M., Jansson, S., and Niyogi, K.K. (2000). A pigment-binding protein essential for regulation of photosynthetic light harvesting. Nature 403, 391–395.

33. Liu, Z., Yan, H., Wang, K., Kuang, T., Zhang, J., Gui, L., An, X., and Chang, W. (2004). Crystal structure of spinach major light-harvesting complex at 2.72 A resolution. Nature 428, 287–292.

34. Maxwell, K., and Johnson, G.N. (2000). Chlorophyll fluorescence - a practical guide. J. Exp. Bot. 51, 659–668.

35. Mendes-Pinto, M.M., Sansiaume, E., Hashimoto, H., Pascal, A.A., Gall, A., and Robert, B. (2013). Electronic absorption and ground state structure of carotenoid molecules. J. Phys. Chem. B 117, 11015–11021.

36. Naranjo, B., Mignee, C., Krieger-Liszkay, A., Hornero-Mendez, D., Gallardo-Guerrero, L., Cejudo, F.J., and Lindahl, M. (2016). The chloroplast NADPH thioredoxin reductase C, NTRC, controls non-photochemical quenching of light energy and photosynthetic electron transport in Arabidopsis. Plant Cell Environ. 39, 804–822.

37. Nilkens, M., Kress, E., Lambrev, P., Miloslavina, Y., Muller, M., Holzwarth, A.R., and Jahns, P. (2010). Identification of a slowly inducible zeaxanthin-dependent component of non-photochemical quenching of chlorophyll fluorescence generated under steady-state conditions in Arabidopsis. Bba-Bioenergetics 1797, 466–475.

38. Niyogi, K.K., Grossman, A.R., and Bjorkman, O. (1998). Arabidopsis mutants define a central role for the xanthophyll cycle in the regulation of photosynthetic energy conversion. Plant Cell 10, 1121–1134.

39. Park, S., Fischer, A.L., Li, Z.R., Bassi, R., Niyogi, K.K., and Fleming, G.R. (2017). Snapshot transient absorption spectroscopy of carotenoid radical cations in high-light-acclimating thylakoid membranes. J. Phys. Chem. Lett. 8, 5548–5554.

40. Park, S., Steen, C.J., Lyska, D., Fischer, A.L., Endelman, B., Iwai, M., Niyogi, K.K., and Fleming, G.R. (2019). Chlorophyll-carotenoid excitation energy transfer and charge transfer in Nannochloropsis oceanica for the regulation of photosynthesis. Proc. Natl. Acad. Sci. USA 116, 3385–3390.

41. Pawlak, K., Paul, S., Liu, C., Reus, M., Yang, C.H., and Holzwarth, A.R. (2020). On the PsbS-induced quenching in the plant major light-harvesting complex LHCII studied in proteoliposomes. Photosynth Res 144, 195–208.

42. Ruban, A.V. (2016). Nonphotochemical chlorophyll cluorescence quenching: Mechanism and effectiveness in protecting plants from photodamage. Plant Physiol. 170, 1903–1916.

43. Ruban, A.V., and Wilson, S. (2020). The Mechanism of Non-photochemical Quenching in Plants: Localisation and Driving Forces. Plant Cell Physiol.

44. Ruban, A.V., Philip, D., Young, A.J., and Horton, P. (1997). Carotenoid-dependent oligomerization of the major chlorophyll *a*/*b* light harvesting complex of Photosystem II of plants. Biochemistry 36, 7855–7859.

45. Ruban, A.V., Pascal, A.A., Robert, B., and Horton, P. (2001). Configuration and dynamics of xanthophylls in light-harvesting antennae of higher plants. Spectroscopic analysis of isolated light-harvesting complex of photosystem II and thylakoid membranes. J. Biol. Chem. 276, 24862–24870.

46. Ruban, A.V., Berera, R., Ilioaia, C., van Stokkum, I.H., Kennis, J.T., Pascal, A.A., van Amerongen, H., Robert, B., Horton, P., and van Grondelle, R. (2007). Identification of a mechanism of photoprotective energy dissipation in higher plants. Nature 450, 575–578.

47. Sacharz, J., Giovagnetti, V., Ungerer, P., Mastroianni, G., and Ruban, A.V. (2017). The xanthophyll cycle affects reversible interactions between PsbS and light-harvesting complex II to control non-photochemical quenching. Nat. Plants 3.

48. Sek, A., Welc, R., Mendes-Pinto, M.M., Reszczynska, E., Grudzinski, W., Luchowski, R., and Gruszecki, W.I. (2020). Raman spectroscopy analysis of molecular configuration forms of the macular xanthophylls. J. Raman Spectr. in press.

49. Siefermann-Harms, D. (1984). Evidence for a heterogenous organization of violaxanthin in thylakoid membranes. Photochem. Photobiol. 40, 507–512.

50. Siefermann, D., and Yamamoto, H.Y. (1974). Light-induced de-epoxidation of violaxanthin in lettuce chloroplasts. III. Reaction kinetics and effect of light intensity on de-epoxidase activity and substrate availability. Biochim. Biophys. Acta 357, 144–150.

51. Son, M., Pinnola, A., and Schlau-Cohen, G.S. (2020a). Zeaxanthin independence of photophysics in light-harvesting complex II in a membrane environment. Biochim. Biophys. Acta 148115, doi: 10.1016/j.bbabio.2019.148115.

52. Son, M., Pinnola, A., Gordon, S.C., Bassi, R., and Schlau-Cohen, G.S. (2020b). Observation of dissipative chlorophyll-to-carotenoid energy transfer in light-harvesting complex II in membrane nanodiscs. Nat Commun 11, 1295.

53. Standfuss, R., van Scheltinga, A.C.T., Lamborghini, M., and Kuhlbrandt, W. (2005). Mechanisms of photoprotection and nonphotochemical quenching in pea light-harvesting complex at 2.5A resolution. EMBO J. 24, 919–928.

54. Steen, C.J., Morris, J.M., Short, A.H., Niyogi, K.K., and Fleming, G.R. (2020). Complex Roles of PsbS and Xanthophylls in the Regulation of Nonphotochemical Quenching in Arabidopsis thaliana under Fluctuating Light. J Phys Chem B 124, 10311–10325.

55. Sylak-Glassman, E.J., Malnoe, A., De Re, E., Brooks, M.D., Fischer, A.L., Niyogi, K.K., and Fleming, G.R. (2014). Distinct roles of the photosystem II protein PsbS and zeaxanthin in the regulation of light harvesting in plants revealed by fluorescence lifetime snapshots. Proc. Natl. Acad. Sci. USA 111, 17498–17503.

56. Tamm, L.K., and Tatulian, S.A. (1997). Infrared spectroscopy of proteins and peptides in lipid bilayers. Q. Rev. Biophys. 30, 365–429.

57. Toth, T.N., Rai, N., Solymosi, K., Zsiros, O., Schroder, W.P., Garab, G., van Amerongen, H., Horton, P., and Kovacs, L. (2016). Fingerprinting the macro-organisation of pigment-protein complexes in plant thylakoid membranes in vivo by circular-dichroism spectroscopy. Biochim. Biophys. Acta 1857, 1479–1489.

58. Wehner, A., Grasses, T., and Jahns, P. (2006). De-epoxidation of violaxanthin in the minor antenna proteins of photosystem II, LHCB4, LHCB5, and LHCB6. J. Biol. Chem. 281, 21924–21933.

59. Welc, R., Luchowski, R., Grudzinski, W., Puzio, M., Sowinski, K., and Gruszecki, W.I. (2016). A key role of xanthophylls that are not embedded in proteins in regulation of the photosynthetic antenna function in plants, revealed by monomolecular layer studies. J. Phys. Chem. B 120, 13056–13064.

60. Wilk, L., Grunwald, M., Liao, P.N., Walla, P.J., and Kuhlbrandt, W. (2013). Direct interaction of the major light-harvesting complex II and PsbS in nonphotochemical quenching. Proc. Natl. Acad. Sci. USA 110, 5452–5456.

61. Xu, P.Q., Tian, L.J., Kloz, M., and Croce, R. (2015). Molecular insights into Zeaxanthin-dependent quenching in higher plants. Sci. Rep. 5, 13679.

62. Yamamoto, H.Y., and Higashi, R.M. (1978). Violaxanthin de-epoxidase - lipid-composition and substrate-specificity. Arch. Biochem. Biophys. 190, 514–522.

63. Zhou, J., Sekatskii, S., Welc, R., Dietler, G., and Gruszecki, W.I. (2020). The role of xanthophylls in the supramolecular organization of the photosynthetic complex LHCII in lipid membranes studied by highresolution imaging and nanospectroscopy. Biochim. Biophys. Acta 1861, 148117.

